# Combinatorial screening of biological degrader modules identifies heterobifunctional molecules that efficiently destroy MYC, leading to rapid tumour cell death *in vitro* and *in vivo*

**DOI:** 10.64898/2026.07.16.738859

**Authors:** Camilla Ascanelli, Samuel Gilberto, Christopher Batho, Gülesin Tunali, Min Ma, Matthew Glover, Adam Seaton, Jonathan Taylor, Hannah Richmond, Richard Butler, Sergio Cuesta Martinez, Sonja Hess, Walid Khaled, Ralph Minter, Heike Laman, Laura Itzhaki, James Hunt, Catherine Wilson

## Abstract

Targeted protein degradation is a potent strategy against intracellular proteins impervious to traditional drugs. We describe a platform reliant on high-throughput cloning and mRNA-based delivery to quickly screen 100s-1000s of modular biological degraders, which we challenge against the high-turnover c-Myc oncoprotein. We uncover critical principles to drive degrader discovery against any target, free from tagging or prior cell line engineering, with direct applications in research and potentially therapy. We demonstrate the feasibility of targeted protein degradation of c-Myc across in vitro cancer models and in vivo xenograft model, uncovering distinct cellular responses between c-Myc degradation and inhibition.

## Main

The disruption of protein function is central to therapy and dissection of biological phenomena. Breakthrough targeted protein degradation (TPD) technologies permit acute protein elimination, usually by hijacking the cellular ubiquitin-proteasome system (UPS)^1^. Proteolysis-targeting chimeras (PROTACs) are heterobifunctional drugs that concomitantly recruit a target and amenable E3 ubiquitin ligase to induce ubiquitylation and degradation of the former and typically outperform their small-molecule drug constituents. Because only target recruitment is required (and not direct modulation), PROTACs are well positioned to broaden the “druggable” target space, but their small-molecule nature restricts usage to targets containing ligand-binding pockets^2^. A valuable alternative are genetically-encoded protein fusions analogous to PROTACs (hereafter bioPROTACs) capable of complete, swift and specific endogenous target destruction^3,4^. Their biological nature enables the recruitment of otherwise non-ligandable targets such as intrinsically disordered proteins (IDPs) or even specific protein variants and states to the vicinity of an unrestricted list of E2 and E3 ligases. bioPROTACs are therefore better suited to extend the current limits of drug target space, thrillingly demonstrated in the destruction of difficult-to-drug oncogenic Ras family proteins^5–7^. Still, their implementation to a wider range of targets suffers from a lack of maturity in the field that has only surveyed a narrow fraction of E3 ligases, often destroying GFP fusions instead of endogenous proteins and proof-of-principle studies focused on long-lived proteins more sensitive to stability changes^3,8–10^. Furthermore, a confounding interplay between bioPROTAC domains can limit protein destruction when compatible partners are combined in inefficient configurations^5^, perhaps explaining why target engagement is not itself predictive of degradation^11^.

Degrader development campaigns lack a screening technology that systematically examines bioPROTAC design features to find productive module pairs. We met this limitation by establishing a scalable automated workflow to quickly assemble numerous bioPROTAC domain combinations and measure their cellular effect in an mRNA-based screening approach. We validated our method against a most difficult protein: the transcription factor c-Myc (hereafter MYC), a long sought-after oncology therapeutic target impervious to traditional drug discovery due to its intrinsically disordered protein (IDP) nature. MYC protein has a very short half-life (∼30 min), which has raised scepticism as to whether the protein is amenable for TPD^12–14^. We chose MYC as the ideal target to challenge our screening strategy and to gain critical insight and guide future campaigns against other difficult endogenous proteins. Importantly, we observed distinct cellular consequences of MYC degradation when compared to inhibition, demonstrating that MYC degradation holds high therapeutic potential.

## Results

### Development of an arrayed bioPROTAC screening strategy

BioPROTACs were engineered with a degradation domain (an E3 ligase’s functional region or E3-recruiting moiety, hereafter “DD”) and a target-binding moiety, interspaced by a linker. By design, target binders replace natural E3 ligase substrate-recruitment motifs, and a nuclear localization signal (NLS) was introduced to enhance MYC co-localisation (Figure 1a). To develop MYC degraders, we first established a DNA fragment toolbox encoding for manifold bioPROTAC modules. Specifically, 45 DDs were selected, 11 sourced from the literature (e.g. SPOP^6^ and TRIM21^15^) and 26 new designs intended at exploring several E3 ligase families and distinct E3 recruitment types to identify optimal attributes (Supplementary Figure 1a). All DDs were chosen based on their suitability for employment in bioPROTACs and the likelihood of encountering MYC (Supplementary Table 1), and new designs partly centred on bacterial/viral proteins to leverage their qualities in UPS hijacking^16^. For MYC recruitment, we sourced two VH single-domain antibody fragments from the literature^17^, CMYCVH-12-321 and CMYCVH-15-321 (hereafter VH12 and VH15, respectively), and previously developed DARPins (DPs) DP07 and DP20 in-house by phage display^18^. All binders were originally identified as standalone inhibitors. Intracellular binder-MYC engagement was confirmed by co-immunoprecipitation in HCT116 cells (Supplementary Figure 1b). Finally, because the influence of linkers in upholding favourable bioPROTAC structural arrangements is yet to be addressed, we included 5 and 16 amino acid residues (AA) glycine-serine flexible linkers, a long 65AA rigid linker^19^, and an 81AA rigid/flexible combination (named L1 to L4, respectively).

**Figure 1:**
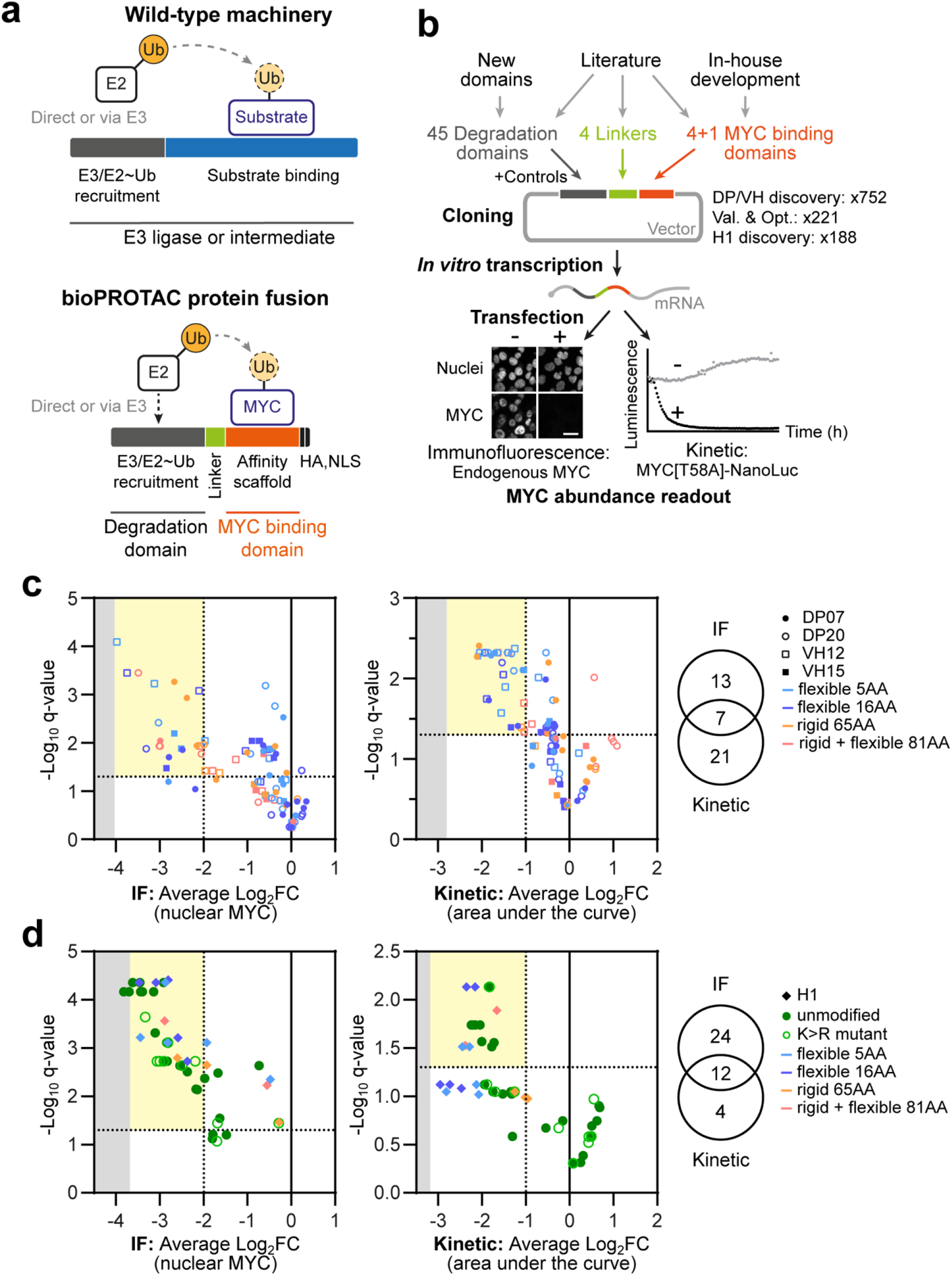
Development of an arrayed bioPROTAC screening strategy pinpoints module combinations capable of downregulating MYC. (**a**) bioPROTACs are constructed by extracting functional domains from E3 ligases or E3 ligase-recruiting proteins, which are fused to a MYC-recruiting scaffold. The E3 remnant (“degradation domain”, DD) must retain the capacity to recruit loaded E2 ubiquitin-conjugating enzymes (E2∼Ub) to facilitate the transfer of ubiquitin to the target. Two specific engineering examples are given for TRIM21^15^ and SPOP^6^-based bioPROTACs (an E3 ligase and intermediate, respectively). (**b**) Workflow for the cloning and screening of manifold fusions. The number of genetic assemblies includes assembled controls. MYC abundance was measured using the indicated methods after mRNA-based bioPROTAC expression in HCT116 cells. Val. & Opt.: Validation & optimization. Scale bar: 25 µm. (**c**, **d**) Confirmatory screening results (average of 3 independent experiments) for the (**c**) DP/VH discovery screening, with bioPROTACs containing DP07, DP20, VH12 or VH15 or (**d**) H1 discovery screening with bioPROTACs containing H1 or the concomitant validation & optimization run with unmodified or K>R mutant bioPROTACs. In discovery runs, colours identify the linker: flexible 5 amino acids (AA), flexible 16AA, rigid 65AA, rigid+flexible 81AA. Volcano plots for 17h end-point immunofluorescence (left) or 48h kinetic data (middle) incorporate average Log_2_FC nuclear MYC or area under the curve for bioPROTAC-expressing samples vs matching control samples. Significance was evaluated by false discovery rate (FDR)-controlled t-tests. FC and q-values threshold is represented by dotted lines, and shaded areas indicate the detection limit (IF) or most extensive MYC downregulation attainable (kinetic assay), determined by cycloheximide treatment. Venn diagrams (right) depict top-performing constructs (Log_2_FC<-2, q-values<0.05 for IF readout and Log_2_FC<-1, q-values<0.05 in kinetic assay).

We established a miniaturised automated workflow to quickly assemble and test all possible toolbox module combinations as an array (“DP/VH discovery screening”), centred on Golden Gate cloning and *in vitro* transcription (Figure 1b). Decisive protocol simplifications were introduced to enable automation without noticeably compromising fidelity (Supplementary Figure 2a), including avoiding plasmid clonal selection, sequencing or mRNA quantification (instead transfected in saturating amounts). Strikingly, mean HA intensity indicated ∼100% of cells exhibited construct expression when using this method (Supplementary Figure 2b). Endogenous MYC abundance changes were measured by immunofluorescence (IF) or kinetically monitored by luminescence in cells expressing NanoLuc-tagged MYC[T58A]. The T58A mutation increases MYC half-life to ∼80 min^20^ and was chosen to enhance degrader selection over endogenous MYC. 720 bioPROTACs were assessed in a primary screening without biological replicates (Supplementary Figure 2c and Supplementary Table 3), and a subset (91) was selected for follow-up confirmation experiments, based on effect (z-score<-1.5 in either assay, avoiding L1/L2 and L3/L4 linker redundancy) or for design feature dissection. We identified 7 strong bioPROTACs (q-values<0.05, Log_2_fold change (FC)<-2 in IF/Log_2_FC<-1 in the kinetic assay), with different combinations of DDs and MYC binding domains, underscoring the need for such a combinatorial approach (Figure 1c). Notably, we observed a significant correlation between bioPROTAC abundance and effect (Pearson’s r=-0.63, p<0.0001) (Supplementary Figure 2d).

Given that E3 ligases can be co-depleted during TPD^21^, we hypothesised that some potent bioPROTACs are destabilised by self-ubiquitylation. Thus, we integrated rationally-designed lysine-arginine substitutions (KR) in hits and low-abundant constructs (compositional redundance was avoided) in an ensuing screening run. We also introduced new non-MYC-binding controls to further validate the hits (thus termed “validation & optimisation run”), and performed in parallel a new discovery run using the original DD/Linker toolbox coupled to another MYC binder, the H1^S6A,F8A^ peptide^22^ (hereafter H1) to further increase compositional diversity. Primary and confirmatory runs were conducted as above, yielding 12 strong degraders common to both assays (Figure 1d, Supplementary Figure 3 and Supplementary Table 4). An attempt to reduce ICP0 domain length failed (Supplementary Figure 3b). KR substitutions enhanced bioPROTAC abundance and effect (Supplementary Figure 4a-c), though most changes were subtle except for the dramatic improvement of DP20-L1-SPOP (Supplementary Figure 4c).

We further extracted favourable bioPROTAC features concealed in our datasets and corroborated specific characteristics by immunoblotting. We noted that 1) the strongest DDs incorporate RING-family E3 ligases, particularly ICP0; 2) module relationships are intricate and unpredictable, as demonstrated by DD-specific binder preferences (Supplementary Figure 5a); 3) all binders can assemble proficient degraders if combined with the right DD, thus finding the right domain pair likely supersedes affinity properties; 4) better degradation effects can be achieved by changing linkers or orientations, as seen for ICP0 (Supplementary Figure 5b); and 5) domain selection govern bioPROTAC efficacy seemingly by influencing both abundance and catalytic properties.

We have previously shown that E2 ubiquitin-conjugating enzymes can be successfully employed to degrade key intracellular oncogenic targets SHP2 and KRAS^7^, therefore we sought to address whether replacing the E3-based DD with UBE2B (hereafter E2B) and UBE2D1 (hereafter E2D1) would successfully degrade nuclear MYC. We generated a panel of twelve E2 bioPROTACs by fusing either E2B or E2D1 to one of the three MYC-binders validated in the E3-based bioPROTACs: VH15^1KR^, DP20^7KR^, or H1, and retaining the linker preference of the original binder. Each construct was tested in two orientations (Binder-E2 and E2-Binder), since the relative position of the MYC-binder to the E2 enzyme may influence substrate recruitment and ubiquitin transfer efficiency due to steric or spatial effects. Construct expression was first validated via *in vitro* translation using the rabbit reticulocyte lysate system, with resulting proteins probed by Western blotting. Eleven of the 12 E2-based bioPROTACs efficiently translated, with the proteins matching the expected molecular weights (Supplementary Figure 6a). Assessing their expression and subcellular localisation in A549 cells, all 11 E2-bioPROTACs constructs demonstrated HA expression and 9 showed successful expression in the nucleus; however, the N-terminal positioning of the H1 peptide disrupted nuclear localisation of the bioPROTACs both when fused to E2D1 or to E2B (Supplementary Figure 6b). Single-cell quantification of MYC levels in bioPROTAC (HA)-expressing cells revealed that E2 bioPROTACs do not significantly deplete MYC at 17 or 24h post-transfection. Notably, E3-based constructs VH15^1KR^-L1-ICP0^1KR^, DP20^7KR^-L1-SPOP and TRIM21-L2-H1 demonstrated evident peak activity at 17h, with a rebound in MYC-positive bioPROTAC-expressing cells at 24h (Supplementary Figure 6c-d). In support of our previous findings, constructs containing the E2B enzyme showed consistently low expression levels compared to E2D1-based constructs (Supplementary Figure 6e). We hypothesised that E2-based degraders might be subject to high turnover rates, possibly due to auto-ubiquitination, limiting their expression and efficacy. Indeed, pulse treatment with proteasome inhibitor MG132 (10 µM) for 1h prior to fixation revealed significant increases in the percentage of cells expressing both E2B-L1-DP20^7KR^ and E2D1-L1-DP20^7KR^ but not the E3-based DP20^7KR^-L1-SPOP or non-degrader control (Supplementary Figure 6f). While limited, our E2-based screen reinforced our previous conclusions on bioPROTAC design, whereby construct orientation needs to be tested in multiple configurations, as it can affect cellular localisation as well as target degradation. In addition, DD-binder preference was again demonstrated, as none of the previously successful binders were able to lead to significant MYC reduction when paired to E2s. Finally, the low protein stability of E2-based constructs, likely due to auto-ubiquitination, is limiting especially when targeting unstable, rapidly turned-over proteins like MYC.

### Validation of MYC-targeting bioPROTAC specificity

Three E3-based bioPROTACs without any domain overlap were chosen for further scrutiny (Figure 2a): VH15^1KR^-L1-ICP0^1KR^, DP20^7KR^-L1-SPOP and TRIM21-L2-H1. The SPOP-based degrader was chosen as an example of E3 indirect recruitment, which underperforms in the kinetic assay yet excels in endogenous MYC destruction. Densitometry analysis following immunoblotting revealed a large reduction in MYC levels (Figure 2b). Furthermore, we characterized the mechanism-of-action of these degraders by mutating vital sites or altogether removing functional domains. All bioPROTACs require direct MYC binding and intact E3 ligase activity to function (Figure 2b). In the case of DP20^7KR^-L1-SPOP, we additionally depleted the recruited E3 ligase’s subunit CUL3 to substantiate our findings. H1 can itself elicit mild MYC downregulation, which is expected given that its stand-alone role in preventing MYC heterodimerization with its interacting partner MAX^22^ modulates MYC protein levels^23^. Finally, we demonstrate that bioPROTAC functional domains do not operate efficiently when split by the T2A auto-cleavable peptide (Figure 2b). These data support a mechanism whereby the selected bioPROTACs potently induce endogenous MYC degradation by bringing into close proximity the target and an active E3 ligase.

**Figure 2:**
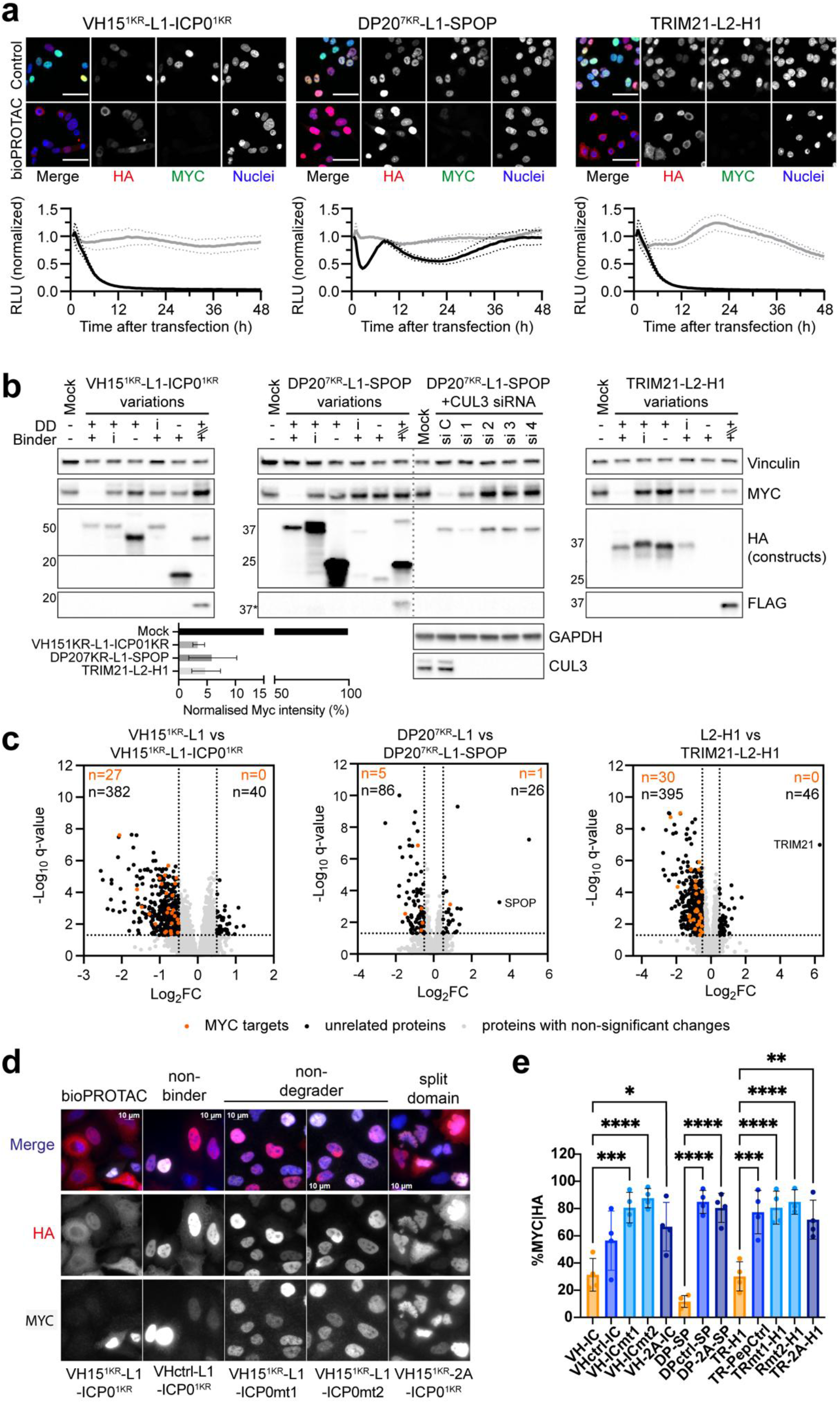
Shortlisted bioPROTACs require concomitant MYC binding and E3 ligase function to eliminate MYC and elicit MYC-specific downstream proteome changes. (**a**) IF (top, scale bar: 50 µm) and kinetic (bottom) MYC abundance readouts for one of n=3 upon expression of the indicated hits from (**a**) the validation and optimisation screening (nonbinding control) and H1 discovery screening (no E3 recruitment control). Time-course data depicts normalized relative light units (RLU) in samples (**–**) and matching controls (**–**). Data is the mean (solid lines) ±standard deviation (SD, dotted lines) of technical replicates in a representative experiment. (**b**) Hit validation by immunoblotting (representative of n=3) after 17h mRNA-based expression of constructs containing (+), lacking (-) or bearing inactive (i) domains, or split by a T2A autocleavable peptide (+\\+) with a resulting N-terminal FLAG tag remnant. The H1 peptide cannot be detected due to its small size. RNAi targeting CUL3 was used 48h in advance to prevent E3 ligase assembly. *molecular weight range corresponds to the unsplit construct. Vinculin and GAPDH serve as loading controls. Densitometry analysis of n=3 independent biological experiments with MYC intensity normalised to Mock. (**c**) Volcano plots depicting proteome changes upon mRNA-mediated expression of the bioPROTACs vs the corresponding unfunctionalized binder controls. Significant changes of protein levels (Log_2_FC≤-0.5 or ≥0.5 and FDR-adjusted p-values (q-values)≤0.001) from 3 independent experiments are depicted for MYC transcriptional targets (orange text) or unrelated proteins (black text); proteins with non-significant changes in levels (grey text). All expressed constructs contain a C-terminal HA tag and NLS. (**d**) Representative images of A549 transfected with VH15^1KR^-L1-ICP0^1KR^ and controls (detected with HA) (scale bar 10 µm). (**e**) Quantification of the IF screen in A549. Data is the percentage of MYC-positive HA-positive nuclei of n=4 biological replicates ± SEM, analysed by One-Way ANOVA. *p<0.05, ** p<0.005, *** 0.0001<p<0.0005, **** p<0.0001

We examined bioPROTAC specificity by measuring global proteome changes by mass spectrometry (Figure 2c, Supplementary Table 5). Construct expression and MYC destruction in these samples was confirmed by immunoblotting (Supplementary Figure 7a). SPOP and TRIM21 were identified in high abundance in the samples transfected with the relative mRNA (Figure 2c, Supplementary Figure 7c).

MYC activates a vast transcriptional programme orchestrating cell growth and proliferation^24^, therefore, using our previously published data^25^ in which RNA sequencing was performed on heart, liver, lung and kidneys from mice 4h following MycER^T2^ activation by 4OHT treatment, we concentrated our analysis on confirmed MYC target modulations common to at least two organs. The representation of MYC targets (Log_2_FC≤-0.5 or ≥0.5 for downregulated or upregulated, respectively; q-values≤0.05) in our data implies that all bioPROTACs interrupt MYC functions (Figure 2c). For analysis of mass spectrometry data, changes in protein abundance were quantified only for proteins with at least two unique peptides. Two MYC peptides, both in the bHLH-LZ domain (C-terminus), were recorded in the proteomics (DQIPELENNEKAPK and THNVLER, Supplementary Figure 7b). Neither peptide was detected in cells transfected with VH15^1KR^-L1-ICP0^1KR^ or TRIM21-L2-H1, in accordance with MYC degradation. While no MYC was detected by immunoblotting with an antibody recognising the N-terminal domain of MYC, when DP20^7KR^-L1-SPOP was transfected, these fragments were significantly higher in abundance in the presence of DP20^7KR^-L1-SPOP compared to its non-binder (DPCtrl-L1-SPOP) and binder-only controls (DP20^7KR^-L1).

Expression of non-binding controls was used to examine DD-driven off-target effects and found to be low (n=14) for the ICP0-based bioPROTACs and included huge proteins such as EP400 which appear in the CRAPome^26^ likely due to peptide probability bias (Supplementary Figure 7c). Overexpression of DPCtrl-L1-SPOP resulted in a total of 83 DEPs, 23 of which were shared with the DP20^7KR^-L1-SPOP bioPROTAC (including SPOP itself; Supplementary Figure 7c). Proteins involved in the response to unfolded protein, like RHBDD1, TMBIM6, TOR1B, appear in the DEPs, however, no GO term was significantly enriched. Transfection with TRIM21-L2-PepCtrl resulted in a total of 226 DEPs, 94 of which were shared with the TRIM21-L2-H1. Significant GO biological process terms associated with these 226 DEPs included proteasomal and Ubiquitin-dependent Protein Catabolic Process, and included many structural subunits of the proteasome, E3 ubiquitin ligase substrate adapters and ubiquitin shuttles such as PSMB7, PSMB4, PSMB5, PSMA1, PSMB3, PSMB1, KLHL18, DCAF13, UBQLN1, UBQLN2, UXT and BAG6, indicating proteasomal perturbation following TRIM21 overexpression.

Known MYC interactor MAX was detected as a single unique peptide (SSAQLQTNYPSSDNSLYTNAK) within the mass spectrometry analysis. MAX peptide abundance (Supplementary Figure 7d) was corroborated by immunoblot (Supplementary Figure 7e) across samples and indicated it was depleted in the presence of the MYC-targeting bioPROTACs VH15^1KR^-L1-ICP0^1KR^ and TRIM21-L2-H1 after mRNA transfection. This was confirmed in doxycycline (dox)-inducible cell lines (Supplementary Figure 7f), where DP20^7KR^-L1-SPOP appeared slow-acting in MAX depletion, indicating protein reduction might not be due to bioPROTAC expression directly.

### bioPROTACs can degrade MYC in multiple cellular contexts

We next sought to confirm the ability of degraders to deplete MYC across cell lines via transfection with mRNA. First, we employed HEK293 cells constitutively expressing LgBiT and with a HiBiT tag knocked-in at the C-terminal position of endogenous, wild-type MYC. Through live cell luminescence recording, we confirmed that transfection with VH15^1KR^-L1-ICP0^1KR^ and TRIM21-L2-H1 resulted in a reduction in MYC-HiBiT signal compared to mock control (Supplementary Figure 8a). Area under the curve (AUC) analysis revealed that TRIM21-L2-H1 was able to significantly reduce MYC-HiBiT levels by 1.8-fold compared to mock (Supplementary Figure 8b). Transient DP20^7KR^-L1-SPOP expression failed to reduce MYC-HiBiT levels, confirming the previous observation in HCT116 overexpressing MYC[T58A]-NanoLuc. It is worth noting that no bioPROTAC resulted in complete depletion of MYC-HiBiT; however, our previous data^27^ demonstrates that even the addition of sublethal dose of cycloheximide failed to completely deplete cells of MYC-HiBiT signal.

Next, we compared the ability of the bioPROTACs to deplete endogenous MYC in two non-small cell lung cancer (NSCLC) cell lines: A549 and NCI-H1373. Cells were transfected with mRNA encoding VH15^1KR^-L1-ICP0^1KR^, DP20^7KR^-L1-SPOP and TRIM21-L2-H1 degraders and their respective controls, with 17h incubation with mRNA. All constructs expressed highly in A549 (Figure 2d, Supplementary Figure 8c), and single-cell analysis of MYC degradation in cells expressing the bioPROTACs or controls (HA signal) revealed that MYC is effectively and significantly depleted by each of the three bioPROTAC constructs (<30% MYC-positive cells), with the specificity of degradation supported by the inactive controls (Figure 2e). NCI-H1373 cells transfection efficiency proved lower than A549 (Supplementary Figure 8d) and none of the bioPROTACs tested resulted in complete depletion of endogenous MYC (Supplementary Figure 8e); however, VH15^1KR^-L1-ICP0^1KR^ and TRIM21-L2-H1 significantly reduced the average MYC intensity by 30% and 50% respectively, when compared to non-degrader or split-domain controls, as measured on a per-cell basis (Supplementary Figure 8f).

We additionally explored the ability of lead bioPROTACs to degrade MYC in Hepatocellular Carcinoma (HCC) cell line Huh7. Constructs displayed variable expression efficiency and bioPROTACs were unable to completely deplete cells of endogenous MYC (Supplementary Figure 8g-h); notably however, DP20^7KR^-L1-SPOP was able to reduce the levels of endogenous MYC by 50% compared to controls (Supplementary Figure 8i). Importantly, endogenous MYC levels vary greatly across the cell lines (Supplementary Figure 8j), with no correlation between MYC levels and successful degradation of MYC by bioPROTACs. Transfection efficiency, construct expression kinetics and availability of co-factors like E2 enzymes might be determinants of bioPROTAC success between cell lines. Taken together, the data suggest that, while the ability of bioPROTACs to completely deplete endogenous MYC varies between cell types, their transient expression leads to MYC degradation in all five cell types tested (A549, HCT116, HEK293 MYC-HiBiT LgBiT, NCI-H1373, Huh7).

### MYC degradation and MYC inhibition have distinct cellular consequences

Using engineered HCT116 expressing bioPROTACs or controls under doxycycline transcriptional control, we monitored cell confluence over time and observed no increased density upon bioPROTAC expression (Figure 3a, Supplementary Figure 9a). Cell viability was measured at 72h endpoint by CellTiter-Glo (Figure 3b, Supplementary Table 6). Expression of binding-only VH15^1KR^ and DP20^7KR^ controls leads to a significant reduction in luminescence compared to parental unmodified cells (1.4- and 2.25-fold change, respectively), likely due to their ability to inhibit MYC, comparable with the 2.32-fold reduction in luminescence measured following treatment with MYC inhibitor 10058-F4. All bioPROTAC-expressing cells resulted in an 8.1-, 3.2- and 7.4-fold signal reduction with VH15^1KR^-L1-ICP0^1KR^, DP20^7KR^-L1-SPOP and TRIM21-L2-H1, respectively. Overall, these data argue that VH15^1KR^-L1-ICP0^1KR^ excels in inducing strong MYC destruction and downstream effects (Figure 2c), with limited off targets (Supplementary Figure 7c) compared to the other candidate bioPROTACs. DP20^7KR^-L1-SPOP is less effective but an adequate candidate when MAX co-depletion is a primary concerning point.

**Figure 3:**
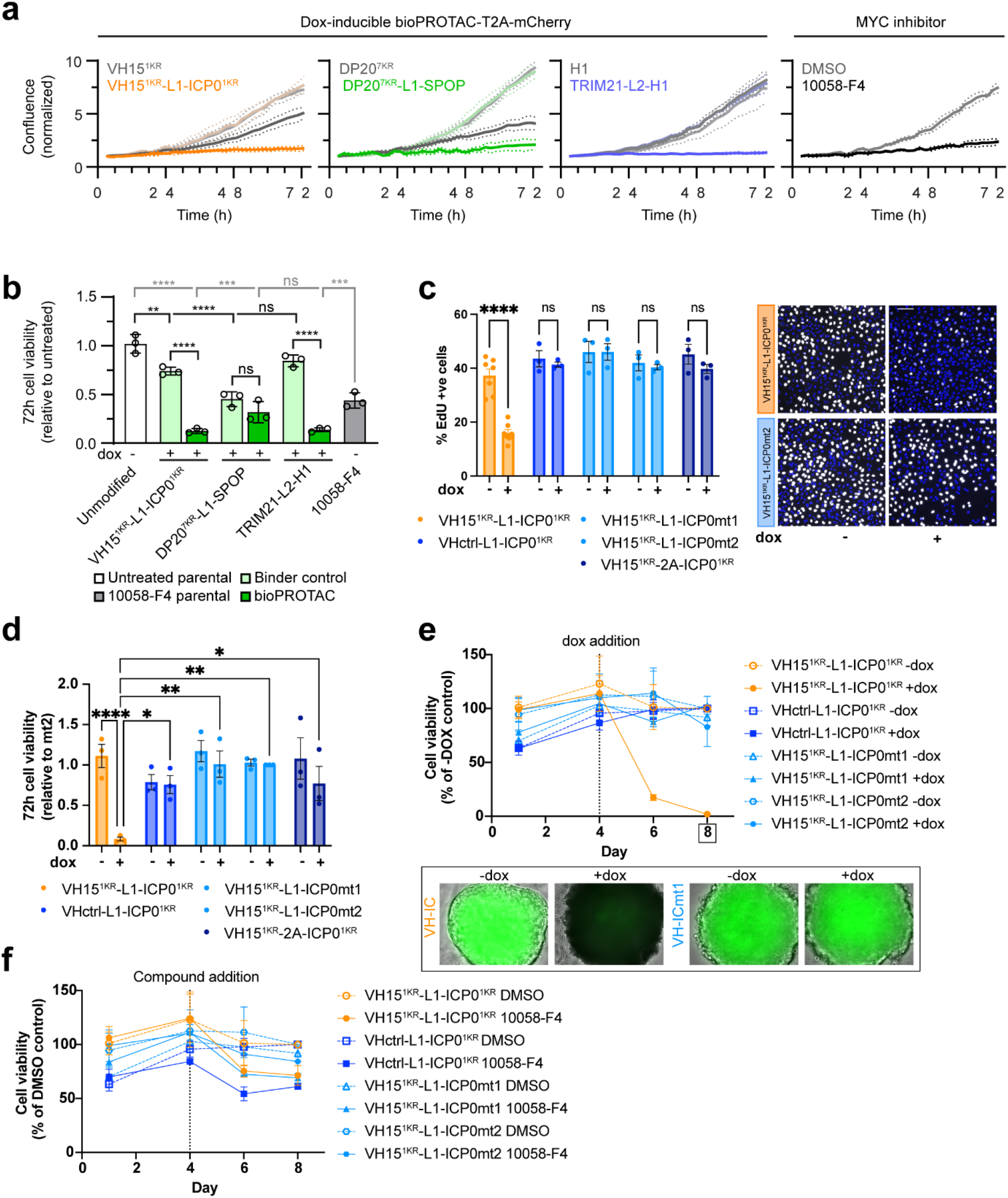
bioPROTAC-mediated MYC degradation leads to distinct cellular consequences to MYC inhibition. (**a**) Live-cell imaging for monitoring of cell confluence (% cell-covered area) of the indicated HCT116 doxycycline-inducible cell lines expressing construct-T2A-mCherry fusions or inhibitor-treated unmodified cells (60 µM 10058-F4). For inducible cell lines, strong colours designate induction of the indicated construct, as opposed to non-induced conditions represented by fainter colours. Continuous lines are the mean of 3 technical replicates for a representative biological replicate (of n=3). Dotted lines depict the error (SD). (**b**) Cell growth was further addressed via a CellTiter Glo 72h end-point assay in the same conditions. Values obtained following induction with doxycycline or treatment with 10058-F4 are depicted relative to untreated controls (no doxycycline or DMSO, respectively). Sample key: parental cells (white), cells expressing dox-inducible binder-only controls (light green) or bioPROTACs (dark green) treated with doxycycline, or parental cells treated with 60 µm 10058-F4 (grey). Shown is the mean of 3 biological replicates (columns) ±SD (error bars). Statistical analysis was performed using a one-way ANOVA followed by Bonferroni multiple comparisons test. (**c**) Cell cycle progression at 24 hours was assessed by EdU incorporation assay in A549 cells expressing doxycycline-inducible VH15^1KR^-L1-ICP0^1KR^ (orange) or non-binder (royal blue), non-degrader (light blue) or split-domain (deep blue) controls. Shown is the mean of 3 biological replicates (columns) ±SEM (error bars), statistical analysis was performed using a two-way ANOVA with multiple comparison test. (scare bar 50 µm) (**d**) Cell viability was assayed in A549 by Crystal Violet staining 72 hours following the induction of expression VH15^1KR^-L1-ICP0^1KR^ (orange) or controls. Data is normalised to VH15^1KR^-L1-ICP0mt2 for n=3 biological replicates (mean ±SEM, two-way ANOVA with multiple comparisons). (**e**) Cell viability of 3D A549 spheroids was assessed by CellTiter Glo over time in cells expressing VH15^1KR^-L1-ICP0^1KR^ or controls in the presence (solid line) or absence (dotted line) of doxycycline on day 4. Data is shown as mean ±SEM (error bars) of n=3 biological replicates. Representative images of spheroids of cells expressing VH15^1KR^-L1-ICP0^1KR^ or VH15^1KR^-L1-ICP0mt1 on day 8. (**f**) As (e) but cells treated with DMSO (dotted line) or MYC inhibitor 10058-F4 (50 µM, full line) from day 4. p-values: *<0.05, **<0.01, ***<0.001, ****<0.0001, ns=non-significant (>0.05).

Building on the observation that VH15^1KR^-L1-ICP0^1KR^ yielded specific degradation of MYC with little DD-driven off-target effects, we generated A549 cells expressing VH15^1KR^-L1-ICP0^1KR^ or controls under doxycycline transcriptional control using lentiviral constructs (Supplementary Figure 9b-c). Modulation of MYC levels across the population was confirmed by IF, with a significant reduction in MYC-positive nuclei at 24h in cells expressing VH15^1KR^-L1-ICP0^1KR^ (Supplementary Figure 9d) compared to controls. Some reduction in MYC-positive cells was seen in the split-domain control VH15^1KR^-T2A-ICP0^1KR^ (Supplementary Figure 9d), likely due to an inefficient cleavage of the T2A construct, resulting in a full-length functional degrader (Supplementary Figure 9c, upper band in lane 10). Importantly, only degradation of MYC induced by VH15^1KR^-L1-ICP0^1KR^ and not inhibition by 10058-F4 led to a significant reduction in MYC-positive cells and MYC intensity in the engineered cell line (Supplementary Figure 9e-f). To validate that the reduction in MYC intensity was due to proteasomal degradation, cells expressing VH15^1KR^-L1-ICP0^1KR^ were treated for 24h in the presence or absence of doxycycline, and in the presence of proteasome inhibitor MG132 (10μM) for the final 2h of incubation. Analysis of protein lysates reveals a reduction in MYC intensity upon expression of VH15^1KR^-L1-ICP0^1KR^ via doxycycline, which is rescued by proteasomal inhibition (Supplementary Figure 9g). Having validated the cell lines for bioPROTAC-induced MYC degradation, we addressed the cellular consequences of MYC degradation by measuring cell proliferation and viability. IF quantifications of 24h doxycycline-treated cells encoding VH15^1KR^-L1-ICP0^1KR^ or controls revealed a significant 2.3-fold decrease in the rate of S-phase entry upon bioPROTAC activation (Figure 3c), and no significant changes in percentage of mitotic cells (Supplementary Figure 9h). Following 72h expression of VH15^1KR^-L1-ICP0^1KR^, colorimetric analysis of crystal violet intensity revealed a significant reduction in viable cells (15-fold decrease compared to its matched control lacking doxycycline) (Figure 3d). To investigate this further, we performed 3D spheroid viability assays in the A549 stable cell lines. Briefly, cells encoding for VH15^1KR^-L1-ICP0^1KR^ or controls were seeded, and spheroids were left to form for four days before the addition of doxycycline or compounds. Cell viability was measured by CellTiter-Glo at days 1, 4, 6, and 8 post-seeding. Induction of VH15^1KR^-L1-ICP0^1KR^ but not controls resulted in a rapid loss of cell viability, which dropped to 20% within 48h of induction, and no viable cells were observed by 96h of expression (Figure 3e). On the contrary, when the same cells were treated with DMSO or MYC inhibitor 10058-F4 in the absence of doxycycline, no significant cytotoxic effects were observed (Figure 3f).

To corroborate that the loss of viability measured by the colourimetric assay is due to a cytotoxic effect, we evaluated cleavage of Caspase 3 (CC3) as a marker for apoptosis by IF in 2D culture. At 24h post-induction of VH15^1KR^-L1-ICP0^1KR^, a significant induction of apoptosis was observed solely following bioPROTAC expression (15.7-fold increase), not upon treatment with 10058-F4 for 24h (1.8-fold increase, Figure 4a) or upon expression of controls (Supplementary Figure 9i). These data demonstrate that MYC inhibition in A549s and degradation result in two distinct cellular responses.

**Figure 4:**
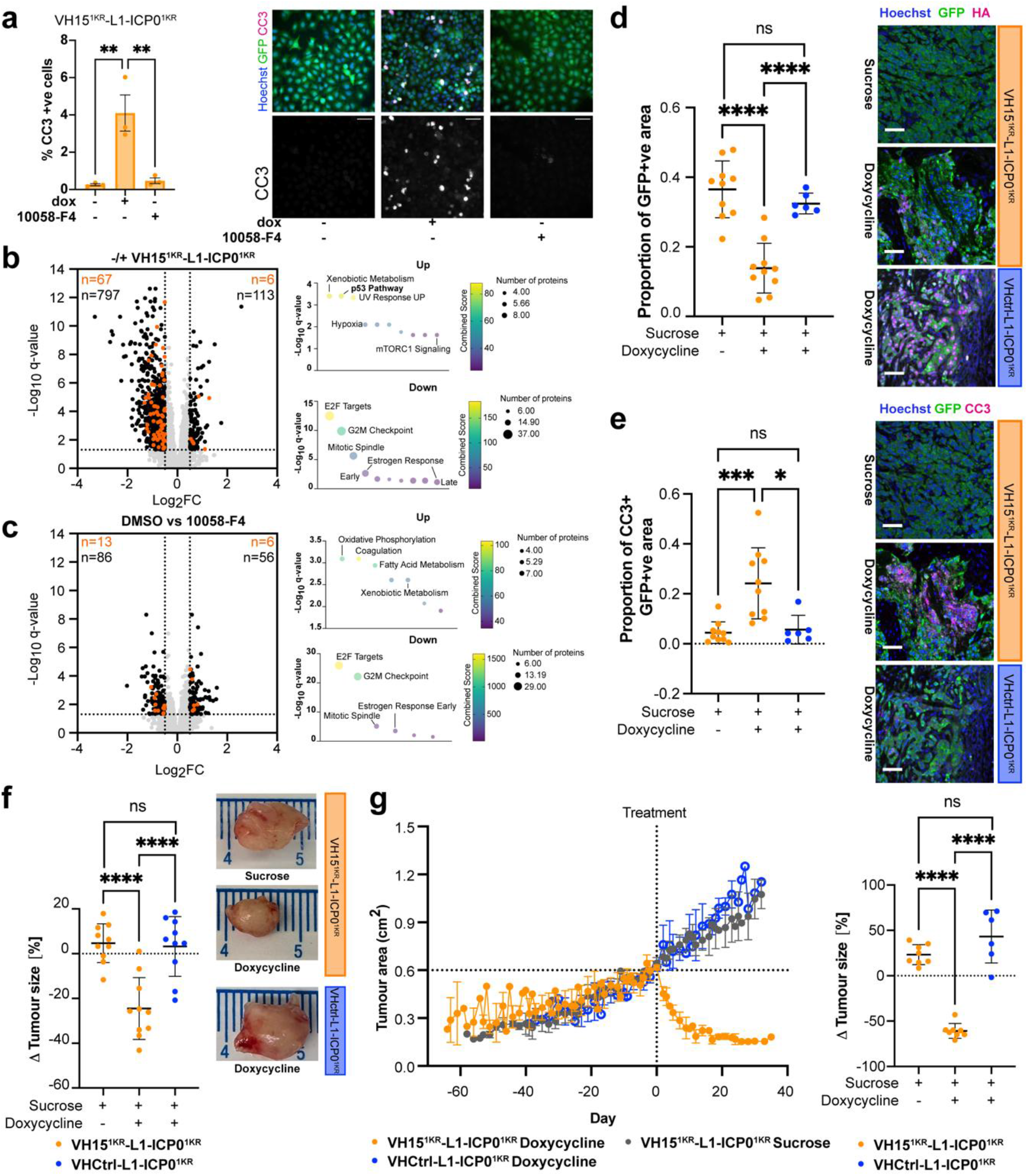
bioPROTAC-mediated MYC degradation leads to acute apoptosis *in vitro* and *in vivo*. (**a**) Induction of cell death by bioPROTACs or MYC inhibitor 10058-F4 (50 µM) was assessed by IF at 24 hours post-treatment with doxycycline or 10058-F4. Shown is the mean of 3 biological replicates (columns) ±SEM (error bars); statistical analysis was performed using Ordinary one-way ANOVA with multiple comparisons. Exemplary immunofluorescence images from one biological replicate probed for Cleaved Caspase 3 (CC3, 50 µm scale bar). Volcano plots depicting proteome changes upon doxycycline expression of VH15^1KR^-L1-ICP0^1KR^ (**b**) or MYC inhibition with 10058-F4 (**c**) vs the corresponding untreated control. Significant changes of protein levels (Log_2_FC≤-0.5 or ≥0.5 and FDR-adjusted p-values (q-values) ≤0.001) from 3 independent experiments are depicted for MYC lung-specific transcriptional targets (orange) or unrelated proteins (black); proteins with non-significant changes in levels (grey). Pathway analysis performed with Enrichr using MSigDB Hallmark (2020) is shown for the corresponding comparisons. (**d**) Histological assessment of A549 xenograft tumours from Athymic nude mice by immunofluorescence demonstrates *in vivo* expression of VH15^1KR^-L1-ICP0^1KR^ or non-binder control VH1ctrl-L1-ICP0^1KR^ (HA) in the doxycycline groups and not the sucrose group (50 µm scale bar). Quantification of GFP-positive sections is shown as Mean ±SD (error bars) with dots representing individual tumours. Statistical analysis was performed by Ordinary one-way ANOVA. (**e**) Induction of apoptosis (Cleaved Caspase 3) upon bioPROTAC expression was confirmed histologically by immunofluorescence (50 µm scale bar) and quantified as a ratio of CC3- to GFP-positive areas. Data shown is Mean ±SD (error bars) with dots representing individual tumours and analysed by ANOVA with Kruskal-Wallis test. Changes in tumour volume 72h (**f**) or 14 days (**g**) post-administration of sucrose or doxycycline. Data is shown as Mean ±SD (error bars) with each dot representing a tumour. Each group contains n=5 mice. (**f**) Tumour growth as measured by tumour area over time for sucrose and doxycycline groups. Data is shown synchronised to treatment start. For VH15^1KR^-L1-ICP0^1KR^ sucrose and doxycycline groups, data are from n=5 mice, VHctrl-L1-ICP0^1KR^ doxycycline n=4 mice. p-values: *<0.05, **<0.01, ***<0.001, ****<0.0001

We next investigated the transcriptional and proteomic consequences of MYC degradation by VH15^1KR^-L1-ICP0^1KR^ in the engineered A549 cell lines via RNA sequencing and mass spectrometry (Supplementary Tables 7-8, respectively). Addition of doxycycline to cells encoding VHctrl-L1-ICP0^1KR^ did not result in any differentially expressed genes (DEGs) or differentially expressed proteins, and a limited number of DEPs (162) were recorded in cells expressing VH15^1KR^-L1-ICP0mt2 (Supplementary Figure 10a-b). In comparison, activation of the bioPROTAC led to over 3000 DEGs and 921 DEPs, while MYC inhibition by 10058-F4 in the same cells resulted in 694 DEGs and 254 DEPs. Upon bioPROTACs activation, we observed a vast overlap in proteomic changes matched in the RNA sequencing data. Specifically, 51.4% of the downregulated DEPs were downregulated at the RNA level, and 61.4% of the upregulated DEPs were also upregulated DEGs (Supplementary Figure 10c-d, respectively), indicating that over half of the protein changes recorded are due to transcriptional changes following MYC degradation and not non-specific degradation by VH15^1KR^-L1-ICP0^1KR^. Next, we addressed the presence of MYC targets among the DEPs in our proteomic data by comparing them with our previously published data set of lung-specific MYC targets^25^. Indeed, 73 lung-specific MYC target genes were found to be significantly changed upon VH15^1KR^-L1-ICP0^1KR^ expression, 67 of which were downregulated, consistent with bioPROTACs-induced MYC degradation (Figure 4b). Only 19 lung-specific MYC-target genes were found to be significantly changed in the same cells treated with 10058-F4 instead of doxycycline (Figure 4c). Pathway analysis with MSigDB Hallmark (2020) revealed similarly downregulated pathways in response to MYC inhibition and degradation (E2F targets, G2M Checkpoint, Mitotic Spindle and Estrogen Response), but distinct upregulated pathways (Figure 4b-c). Treatment with 10058-F4 resulted in upregulation of proteins related to oxidative phosphorylation, coagulation and fatty acid metabolism. Xenobiotic metabolism was noted for both 10058-F4 treatment and activation of VH15^1KR^-L1-ICP0^1KR^ by doxycycline; however, bioPROTACs activation resulted in distinct upregulation of p53, UV response and Hypoxia pathways. Similar pathway changes were recorded in the transcriptomic data, where genes related to the Estrogen Response pathway and E2F targets were significantly reduced in both MYC inhibition and degradation contexts (Supplementary Figure 10e-f), however, MYC degradation by VH15^1KR^-L1-ICP0^1KR^ led only to the induction of p53 pathway genes. Together, our data demonstrate that like MYC inhibition, MYC degradation by bioPROTACs leads to cell cycle exit, however this is uniquely accompanied by an acute and rapid induction of apoptosis.

### MYC- bioPROTAC expression leads to rapid tumour regression *in vivo*

To address the effects of MYC degradation on tumour growth, we conducted xenograft experiments in athymic *NU(NCr)-Foxn1nu* mice. Mice 6 weeks of age received subcutaneous injections in both flanks of (5×10^5^) A549 cells encoding doxycycline-inducible VH15^1KR^-L1-ICP0^1KR^ or VHctrl-L1-ICP0^1KR^. After a period of incubation post-injection to allow tumour establishment, mice were weighed and tumours measured three times per week for the duration of the study. Once tumour size reached 0.6cm^2^, mice received doxycycline water (2g/L doxycycline, 3% sucrose) or sucrose water (3% sucrose). Two cohorts were run in parallel, one longitudinal and one where tissues were collected 72h post administration of doxycycline/sucrose water. Histological analysis of the tumours collected at 72h post-treatment confirmed constitutive GFP expression. Doxycycline addition resulted in expression of bioPROTAC or control constructs (HA staining, Figure 4d, Supplementary Figure 10g). A significant reduction in GFP-positive area in the VH15^1KR^-L1-ICP0^1KR^ doxycycline group compared to both control groups, VH15^1KR^-L1-ICP0^1KR^ sucrose and VHctrl-L1-ICP0^1KR^ doxycycline (Figure 4d) was observed, with 37.6% and 42.8% decrease compared to sucrose VH15^1KR^-L1-ICP0^1KR^ and doxycycline VHCtrl-L1-ICP0^1KR^, respectively. This loss was due to the induction of apoptosis upon bioPROTAC activation alone, as demonstrated by CC3 staining (Figure 4e, Supplementary Figure 10h) with 18% and 23.3% increase compared to Sucrose VH15^1KR^-L1-ICP0^1KR^ and doxycycline VHCtrl-L1-ICP0^1KR^. The cell death was unique to the expression of the bioPROTAC and resulted in a decrease in tumour volume of 24.5% in 72h when compared to sucrose (4.7% increase) and to non-binding VHctrl-L1-ICP0^1KR^ control (3.2% increase) (Figure 4f). Longitudinal studies revealed tumour regression to baseline size within 10 days of treatment. By day 14, administration of doxycycline water in the VH15^1KR^-L1-ICP0^1KR^ group led to a 60.8% reduction in tumour size, but not VH15^1KR^-L1-ICP0^1KR^ sucrose (23.3% increase) and doxycycline VHctrl-L1-ICP0^1KR^ (43.4% increase) control groups (Figure 4g). Overall, these data confirm *in vivo* the acute onset of cell death following bioPROTAC activation seen *in vitro*, and establishes MYC degradation as a powerful modality for rapid tumour regression.

## Discussion

The majority of human cancers feature MYC upregulation, with over half of cancers demonstrating deregulated MYC networks ^28^. MYC is not only an oncogenic driver, but its inactivation triggers tumour regression, underpinning a clear unmet therapeutic opportunity^12,13,24^. Collectively, our study demonstrates that bioPROTACs can be used to directly modulate the high-turnover MYC and provides a roadmap to destroy virtually any endogenous protein (or a target subpopulation) with high specificity, without the need for prior cell line engineering or protein tagging. Accordingly, we elucidate a set of features that characterise exceptional biological degraders and uncover ICP0 as a new and potent DD. Our dataset strongly challenges a one-size-fits-all recipe for bioPROTAC design due to unpredictable domain relationships and, consequently, the traditional stepwise approach for bioPROTAC development must be replaced by combinatorial screening so as not to miss out proficient module pairs. We also expose interactor co-depletion, an effect usually overlooked but which may generally apply to all TPD approaches and should therefore be more routinely investigated. Building on our previous work employing E2 enzymes in bioPROTAC design^7^, we demonstrate adoption of E3-based versus E2-based bioPROTACs likely depends on target selection, localisation and stability. In addition, we reveal that sustained degradation of MYC in A549s leads to a more acute and widespread apoptotic induction to that previously observed with MYC inhibitors^29–32^, *in vitro* and rapid tumour regression in less than two weeks *in vivo* in a xenograft tumour model. We anticipate that with further advancements in gene delivery, this strategy will enable a new generation of therapeutics against MYC and other “undruggable” targets.

## Acknowledgements

We thank the AstraZeneca R&D Postdoc programme for supporting this work. R. Davies for initial project discussions. We also thank our colleagues in the AstraZeneca Nucleic Acid Therapeutics Team for advice on mRNA design, synthesis and delivery; S. Peel for help with microscopy; Biologics Engineering Automation Team for assisting workflow automation; the AstraZeneca Dynamic Omics Team for technical assistance with Proteomics data acquisition. DARPins were discovered under license from Molecular Partners ahead of this work. We are grateful to current and past members of the AstraZeneca Affinity Reagents Team for technical support, discussions and reviewing the manuscript. We apologize to authors whose work couldn’t be cited due to space restrictions. The authors thank the support staff in the Cambridge University Biomedical Services at the Anne McLaren Building and the flow cytometry facility from the School of Biological Sciences for their support and assistance in this work.

## Funding

This work was supported by funding from AstraZeneca (G123209, to C.H.W and L.S.I), Wellcome Trust Developing Concept Fund (G111225 to C.H.W, L.S.I and C.A; G681163 to C.A, C.H.W and L.S.I administered through Cambridge Academy for Therapeutic Sciences), BBSRC (G129853 to C.H.W, L.S.I and Paul Miller).

## Competing interests

This work was supported by AstraZeneca, M.M., M.G., J.T., S.C.M. and J.H. are employees of AstraZeneca and may own AstraZeneca shares. S.G. and R.M. were employees of AstraZeneca while engaged in this research project. S.G. is now an employee of Monte Rosa Therapeutics. R.M. is now an employee of Forth Therapeutics Ltd. J.H. and S.G. are authors on a patent related to this work.

## Author contributions

Conceptualisation (JH, LSI, CHW, HL, CA, SG, RM), Funding acquisition (JH, LSI, CHW, CA), Investigation (CA, SG, CAPB, GT, MG, MM, AWS, HR, SCM), Software (RB), Resources (JH, LSI, CHW, WTK, SH), Project administration (JH, LSI, CHW, WTK), Writing – original draft preparation (CHW, CA, SG, SCM, JH), Writing – review and editing (LSI, HL, JT).

## Sources of funding

This work was supported by funds from AZ, Isaac Newton Trust, Wellcome Trust DCF (Wellcome Institutional Translational Partnership Award 222062/Z/20/Z and G681163 to CA, CHW and LSI) and BBSRC Engineering Biology Seedcorn Award (G123986 to LSI and CHW).

## Data availability

The proteomics dataset generated and analysed during the study is available upon reasonable request.

Raw RNA sequencing data generated in this study are available on Array Express under Accession number E-MTAB-16753. BioPROTAC screening data is available from the corresponding authors upon reasonable request.

## Methods

### Selection and engineering of degradation domains

Selected degradation domains originate from human, bacterial or viral E3 ligases or intermediate proteins capable of recruiting human E3 ligases. Domains were either retrieved from other studies or newly developed, extracted from proteins matching the following criteria: 1) Associated with protein degradation, 2) Functional domains are well understood (biochemically and/or the structure has been determined and is available in the protein data bank (PDB)^33^, www.rcsb.org), ideally continuous and not dependent on post-translational modifications. If the domain is an intermediate, the indirectly-recruited E3 ligase is 3) present in the nucleus (to co-localize with MYC^12^), 4) abundant in a wide range of tissues (to ensure wide applicability) and 5) widely expressed in cancer, ideally overexpressed, for applicability and to ensure enough supply to degrade the overexpressed MYC (sources: Uniprot^34^, www.uniprot.org, Human Protein Atlas^35^, www.proteinatlas.org, and GEPIA^36^, gepia.cancer-pku.cn). Additionally, we favoured intermediates where the recruited partner is found to be essential in >50% of the studies to avoid cellular resistance to the bioPROTACs, though this was not a requirement (source: OGEE v2^37^, currently version 3 v3.ogee.info). We considered an E3 ligase essential if the direct interacting partner is reported as an essential gene in >80% of the studies included in the OGEE v2 database and not essential if in <20% of the studies. For new designs, we compiled a non-exhaustive list of proteins matching the above criteria after manual literature searches, supported by UniProt curated information. To circumvent deleterious gain-of-function, most natural substrate-binding sequences were removed or mutated (e.g. T67A in ICP0^38^), guided by literature observations and UniProt annotations, unless the functional regions cannot be fully separated and a resulting gain-of-function is deemed not harmful. We also explored the benefit of positioning a functional group in either N- or C-terminal relative to the MYC binder when there is evidence that natural substrates may bind in either end, further truncated CRL adaptors βTrCP, FBXO17 and DDB2 in search of a minimal functional domain and tested HUWE1 constructs with and without its autoinhibitory sequence as an attempt of mitigating self-ubiquitylation and degradation. Undesired trafficking sequences are also removed, such as membrane-targeting or nuclear exclusion signals. See Supplementary Table 1.

### Affinity protein generation

α-MYC DARPins 07, 20^18^ and an α-IDOL DARPin were isolated from a phage display library ahead of this work, as previously described^18,39^, by selections against GST-MYC bHLH-LZ (AA 330-439; Novus biologicals) or full-length IDOL. Deselection against GST-Ubiqutin was also performed to eliminate GST-binding molecules. Validation of antigen-DARPin binding was performed by immunoassays as described^39,40^. MDM2 VHHs were generated by Hybrigenics Services SAS, Paris, France via a yeast two-hybrid screening against LexA-MDM2 (AA 1-188). As non-MYC binding scaffold controls, we used DARPin E3_5^41^, a VH targeting hen egg white lysozyme^42^ and an α-helical peptide from the DARPin backbone. Protein sequences of all fragments generated can be found in Supplementary Table 1 and Supplementary Table 2.

### Single insert cloning and mutagenesis

Individual DNA fragments were generated by gene synthesis (Twist Bioscience, GenScript or Invitrogen GeneArt) or PCR, except fragments smaller than 80 bp which were obtained by annealing of two complementary oligonucleotides by heating at 95°C and cooling down to 22°C at a rate of 0.1°C/s. Non-human sequences were subjected to codon optimization (GenScript). DNA encoding for the degradation domains, linkers, MYC-binding domains and controls were cloned into the pCR Blunt II-TOPO vector using the Zero Blunt TOPO PCR Cloning Kit (Thermo Fisher) following the manufacturer’s instructions. The rigid linker was sourced from^19^ (“L3” in the original publication). cDNA encoding for human MYC was subcloned into pFN31K/pFC32K vectors (Promega) for fusion with NanoLuc luciferase (expression controlled by the hPGK promoter), DNA encoding for FLAG-tagged DP07, DP20, DARPin E3_5, VH12 and VH15 were subcloned into the pFC14K vector (Promega), DNA encoding shortlisted bioPROTAC-HA-NLS constructs and controls was subcloned into a modified ODIn-inv-neo mammalian expression plasmid^43^ containing T2A-mCherry. The Golden Gate destination (GG-dest) vector was generated by integration of a cassette comprising the T7 promoter with AG initiator sequence, UTRs^44^, the Kozak consensus sequence, the ccdB killer gene plus a chloramphenicol resistance gene flanked by Esp3I recognition sequences, HA tag and MYC NLS^45^ sequences onto a pcDNA3.1(+) vector (Invitrogen). MYC Thr 58 was mutated to Ala (T58A) following a “round-the-horn” PCR-based protocol. Briefly, after PCR the template vector was digested with DpnI (New England Biolabs, NEB) for 4h at 37°C, purified using the QIAquick PCR purification kit (Qiagen), blunt ends were phosphorylated using PNK (NEB) in 1x T4 ligase buffer for 20 min at 37°C. Following heat-inactivation of PNK at 75°C for 10 min, ends were ligated using T4 ligase (NEB) by incubating at 2h at room temperature and overnight at 16°C. All other mutants were obtained by gene synthesis. PCR was performed using Phusion High-Fidelity Master Mix (NEB) following vendor instructions. Oligonucleotides used for PCR amplification and mutagenesis are listed in Supplementary Table 7. Decisions to mutate lysine residues in degradation domains are based on conservation with orthologs (may include mouse, rat, fruit fly, bovine, pig, zebrafish, western clawed frog and African clawed frog), conservation in human paralogs, presence within critical functional domains and surface exposure (inferred from the positioning of a lysine’s side chain in available tridimensional structures). Mutagenesis of MYC-binding domains relied on the conservation and surface exposure of residues from similar affinity proteins. Sequence and structural information were obtained from UniProt and PDB. See Supplementary Table 1 and Supplementary Table 2. All sequences were analysed using Geneious Prime 2020.

### Automated multi-fragment cloning and *in vitro* transcription

Arrayed unique combinations of toolbox DNA fragments encoding for the degradation domains (or Halo tag as negative control), linkers and MYC-binding domains (or non-binding controls), flanked by Esp3I recognition sequences and pre-cloned into the pCR Blunt II-TOPO vector, were individually assembled by a modified Golden Gate cloning protocol^46^ which resulted in >98% correct clones in preliminary control reactions. Overhang sequences ATGG, GCCG, GAAG and GGAT were chosen based on T4 ligase fidelity information^47^, using the Ligase Fidelity Viewer (NEB; ggtools.neb.com/viewset/run.cgi). 2 μL reactions containing 1 fmol of DNA fragments (3 fragments per reaction), 1 fmol GG-dest vector, 0.1 μL Esp3I FastDigest Enzyme (Thermo Fisher), 0.1 μL T4 DNA ligase 2×10^6^ U/mL (NEB) and 0.2 μL 10x T4 ligase buffer (NEB) were subjected to 30 incubation cycles alternating between 37°C for 5 min and 16°C for 5 min, followed by 30 min at 37°C and 15 min at 75°C. 0.5 μL of a master mix containing 0.125 μL Plasmid-Safe ATP-Dependent DNase (Lucigen/Epicentre) and 0.125 μL ATP 25 mM in 1x T4 DNA ligase buffer (NEB) were then added to each reaction and incubated for 1h at 37°C. High efficiency chemically competent DH5α *E. coli* (NEB) were grown in liquid 2x TY medium overnight at 37°C following transformation, after which 0.5 μL of the liquid culture were directly used for amplification of the T7-5’UTR-bioPROTAC-HA-NLS-3’UTR-polyA cassette by PCR (oligonucleotides in Supplementary Table 7). Following incubation with DpnI for 1h at 37°C, products were bound to AMPure XP beads (Beckman Coulter), washed with 70% ethanol and eluted with RNase-free water. Subsequently, arrayed *in vitro* transcription reactions were performed using the HiScribe T7 high yield RNA synthesis kit (NEB) as per manufacturer instructions, with the following variations: 1) reagent volumes were proportionally reduced for 2 μL reactions, 2) 20% of UTP was replaced with 5-Methoxyuridine-5’-Triphosphate (5-moUTP, TriLink) and 3) co-transcriptional capping was performed by including CleanCap AG (TriLink) in equimolar amount to NTPs. Template DNA was digested with DNase I (NEB). DNA quality control was performed by electrophoresis and Sanger sequencing of a small fraction of PCR products and mRNA integrity and yield was examined using a Bioanalyzer RNA 6000 Nano assay (Agilent) for randomly-selected constructs. mRNA was not purified prior to screening, and the mRNA concentration of screening library elements was extrapolated based on the analysed reaction subset. N.b. we have determined that upon GFP expression, transfection with >50 ng mRNA does increase protein expression levels, i.e. the fluctuation of mRNA yields should not be reflected in bioPROTAC protein abundance following mRNA transfection. For follow-up experiments, plasmid DNA from individual clones was purified and sequenced and PCR reactions were column-purified using a QIAquick PCR purification kit. mRNA was synthesised in 10 μL reactions containing 40% 5-moUTP, purified using a MEGAclear Transcription Clean-Up kit (Invitrogen) or RNeasy 96 Kit (Qiagen) according to vendor instructions and quantified with a Lunatic nucleic acid quantification system (Unchained labs) to allow transfections with equal amounts of mRNA. Instrumentation for screening: purified DNA (plasmids and PCR products) was dispensed by acoustic droplet ejection using an Echo 550 liquid handler (Labcyte). For automated cloning and *in vitro* transcription, shared components (except GG-dest vector) were transferred as a master mix using a Certus Flex liquid dispenser (Fritz Gyger). Other dispensing steps during screening were performed using a Hamilton Microlab Star liquid handling system (Hamilton). DNA sequences were analysed using Geneious Prime 2020.

### Cell culture and cell line generation

All cell culture reagents are from Gibco, Thermo Fisher unless otherwise indicated. All cell lines were maintained at 37 °C and 5% CO_2_. HCT116 cells were purchased from ATCC and cultured in DMEM supplemented with 10% FBS and 1% penicillin–streptomycin. A549 and H1373 were cultured in RPMI supplemented with 10% FBS; HEK293 MYC-HiBiT LgBiT and Huh7 were maintained in DMEM supplemented with 10% FBS; HEK293FT cells for lentiviral generation were maintained in DMEM supplemented with 10% FBS, 0.1mM NEAA, 6mM L-Glutamine, 1mM MEM Sodium Pyruvate. Cell lines with doxycycline-inducible cassettes were maintained in the respective media supplemented with 10%Tetracycline-free FBS. In the absence of a CO_2_-controlled environment during kinetic luminescence experiments, Leibovitz’s L-15 medium was used instead of DMEM. To establish modified cell lines, cells were transfected using FuGene HD (Promega) following manufacturer’s instructions. ObLiGaRe Doxycycline Inducible (ODin)^43^ cell lines were generated by co-transfecting bioPROTAC or control-encoding ODIn-inv-neo and ZFN-AAVS1 plasmids at a ratio 1:2 and selecting with 0.5 mg/mL geneticin-containing growth media. For lentivirally transduced cells, lentivirus were first generated in HEK293FT cells by transfection with PEI MAX (Clinisciences Limited), packaging plasmids PAX2 and PMD2.G, and transfer plasmids encoding for bioPROTAC or controls under Tetracycline control. The backbone for the transfer plasmids, SIN40C.TRE.MCS.IRES.dTomato.PGK.sfGFP.P2A.Tet3G, was a gift from Dirk Heckl (Addgene plasmid #169283 ; http://n2t.net/addgene:169283; RRID:Addgene_169283). The following day, transfection efficiency was evaluated via GFP using an EVOS microscope and if transfection efficiency was above 60%, the media was replaced with destination cell line media for 48-72 hours. Infected media was collected, filtered through a 40mm filter, supplemented with Polybrene (Merk), and placed on destination cells at a 2:1 ratio with fresh media for 72-96 hours before single-cell sorting of GFP on a BD FACSDiscover S8 cell sorter with real-time imaging.

Cells were tested and found to be mycoplasma-free before use.

### Immunofluorescence microscopy

For IF experiments, unmodified HCT116 cells were transfected with mRNA using Lipofectamine RNAiMAX (Invitrogen), which resulted in >99% transfected cells when tested with GFP, as follows: ≥50 ng bioPROTAC-encoding mRNA were added via acoustic dispensing to the wells of 384-well plates containing Opti-MEM (Gibco) in triplicate, onto which 0.15 μL Lipofectamine RNAiMAX in Opti-MEM was added, followed by 15-30 min incubation at room temperature and the seeding of 8000 cells per well. For unmodified A549, NCI-H1373 forward transfection with Lipofectamine RNAiMAX was performed, while reverse transfection was performed in Huh7. Cells were seeded at 25,000-75,000 cells per well (A549 and NCI-H1373, respectively) in a 12 Well Chamber removable slide (ibidi), and transfected 24 hours after seeding with 1.2pmol of mRNA and 0.2 μL of Lipofectamine. 17h post-transfection, cells were incubated in 4% formaldehyde for 15 min, washed 3x with PBS and incubated in blocking buffer (3% BSA, 0.1% Triton X-100 in PBS) for 1h. Fixed cells were incubated overnight with anti-MYC antibody Y69 (Abcam) and anti-HA tag antibody 16B12 (Abcam and Enzo Life Sciences), both diluted 1:5000 in blocking buffer, followed by incubation with Donkey anti-rabbit IgG/Alexa fluor 488 or 647 1:500, Goat anti-mouse IgG/Alexa fluor 568 or 555 1:500 and Hoechst 2 µg/mL (all from Thermo Fisher) in blocking buffer for 1h and 3 washes with PBS. For the screen in HCT116 cells, 4 images per well were acquired by a CV7000 spinning disk confocal microscope (Yokogawa) using a 20x objective. For follow-up screen in A549 and NCI-H1373, 9 images per well were acquired by a Leica DM6B microscope at 10x. Raw images from the Yokogawa were processed using the Columbus 2.9.1.532 software (PerkinElmer) for quantification of nuclear MYC and HA (bioPROTAC) abundance for all cells in the field of view. Raw values per cell are an average of the nuclear pixel intensity for the respective fluorescence channel. The values brought forward per well are the median of the cell population. In each experiment, three technical replicates were measured (these are from distinct samples and not repeated measurements of the same sample). Up to three independent experiments were performed. For A549 and NCI-H1373 IF experiments, a semi-automated image analysis pipeline was set up in CellProfiler. Treatment with 100 μg/mL of the protein synthesis inhibitor cycloheximide (Sigma Aldrich) was used as a proxy for background fluorescence in absence of MYC and provides a measure of the detection limit, based on the estimation of a negligible residual MYC of 5.8×10^-9^ % after 17h treatment (considering a half-life of 30 min and an exponential decay of its abundance). Figure panels were constructed using ImageJ 2.0.0-rc-69/1.52p and Adobe Illustrator 2025 (version 29.7.1). Prior to this study, we confirmed the high specificity of the anti-MYC Y69 antibody by siRNA-mediated downregulation of MYC and that the anti-HA antibody does not yield detectable signal when HA-tagged proteins are not expressed.

### Luminescence kinetic assay

HCT116 MYC[T58A]-NanoLuc-expressing cells were transfected in triplicate following the protocol indicated for the HCT116 IF experiments, except that Nano-Glo Vivazine live cell NanoLuc substrate (Promega) was pre-mixed with cells prior to seeding. Readings were made every 30 min for 48h using an EnVision plate reader (PerkinElmer) with ultrasensitive luminescence detection. In each experiment, three technical replicates were measured (these are from distinct samples and not repeated measurements of the same sample). Up to three independent experiments were performed. Time resolution was lower in primary screens due to practical reasons. 100 μg/mL Cycloheximide treatment was used to estimate the maximum attainable MYC downregulation, based on the assumption that its decay in complete absence of protein synthesis (80 min half-life^20^) is faster than induced protein degradation following mRNA transfection. Residual MYC[T58A] is 3.8×10^-4^ % 24h post-transfection considering a 80 min half-life. For HEK293 MYC-HiBiT LgBit experiments, reverse transfections with Lipofectamine RNAiMAX were performed. Briefly, 1.2pmol of mRNA was incubated with Opti-MEM and 0.2 μL Lipofectamine for 20 minutes, following manufacturer’s instructions, after which 40,000 cells were seeded per well in triplicate in media containing Nano-Glo Endurazine live cell NanoLuc substrate (Promega) premixed prior to seeding. Readings were made every 49 min for 72h using a CLARIOstar Plus luminometer (BMG Labtech).

### Screening data processing

For the IF readout, the median of nuclear MYC measurements for the cell population was normalized to the plate-specific mock control to eliminate plate-to-plate variations and signal disparities across biological replicates. Nuclear HA (bioPROTACs) lacks a universal standard reference, and thus the data was not transformed except for the visualization of bioPROTAC expression relative to the average of controls in Supplementary Figure 2 and Supplementary Figure 4. HA measurements were therefore not corrected for plate-to-plate variations. In the kinetic assay, luminescence relative light unit (RLU) values were normalized to the respective mock control data points to eliminate time-point- and plate-specific variations. Kinetic datasets were then normalized to the first measured data point, and the area under each curve was calculated following the trapezoidal rule. Values for technical replicates were averaged and nuclear MYC (IF) or area under the curve (kinetic) fold changes (FC) were calculated as the MYC abundance ratio in the presence of bioPROTACs vs specific innocuous control. For discovery-type screenings, controls contain the linker and MYC binder, but lack a degradation domain (replaced by the Halo tag) and for the validation & optimization screening, controls use target-binding scaffolds which do not recruit MYC. In the primary screenings, FCs were standardized as z-scores to evaluate performance relative to the dataset samples, also providing a dimension to compare both readouts. Selected hits were carried forward for confirmation by performing two additional independent experiments (i.e. final n=3). Wells with median HA values (in IF) lower than the average of mock samples’ medians + 3x standard deviation (SD) were considered expression-, transfection- or staining-negative and were excluded on grounds of evidence of technical error. Accordingly, MYC abundance readings in the IF and kinetic assays from constructs not detected in any technical replicate were excluded, thereby resulting in some cases with 2 usable biological replicates, as indicated (Supplementary Table 3 and Supplementary Table 4). Where a control was not detected in primary screenings, the most similar one was chosen to calculate FCs (does not apply to confirmatory runs). Statistical analysis of MYC abundance and area under the curve measurements were performed vs sample-specific controls using multiple unpaired two-tailed t-tests corrected for FDR with the two-stage Benjamini, Krieger and Yekutieli method. We assumed a normal distribution of the biological replicate values, and that a given sample has the same SD as its respective control (i.e. we didn’t assume that all SDs are equal). Reported are Log_2_FC, difference between mean values and associated standard error, t ratio (difference between means divided by the standard error of the difference), degrees of freedom and q-values (FDR-adjusted p-values). Discoveries were assigned based on a 5% FDR threshold and Log_2_FC < -2 (IF) or <-1 (Kinetic assay). bioPROTAC expression and MYC abundance correlations were evaluated by Pearson correlation tests, with reported correlation coefficient (r) and two-sided p-value. Basic calculations and data handling were performed in MS Excel (v2102) and all statistical analyses were performed using the GraphPad Prism 9 software. For analysis of MYC degradation in HEK293 MYC-HiBiT LgBit, for each biological replicate, raw luminescence data of each condition was normalised to the average mock luminesce, followed by normalisation to time 0, generating FC in raw luminescence units (RLU). Per biological replicate, Average FC in RLU for three technical replicates was plotted, and AUC calculations were performed on GraphPad Prism software. For analysis of MYC degradation in A549, NCI-H1373, a CellProfiler pipeline was set up, identifying three distinct primary objects (nuclei, HA objects, MYC objects) and related to define HA-positive + MYC-positive nuclei. The percentage of HA-positive nuclei also positive for HA was calculated, and the average of four biological replicates was plotted and analysed by One-way ANOVA. When MYC intensity was analysed, MYC intensity for HA-positive nuclei was measured via CellProfiler and normalised to the intensity of a non-degrading control. For analysis of S-phase rate, the pipeline was adapted to identify EdU-positive objects, which were related and filtered to the nuclei detected by Hoechst staining. Percentage of EdU-positive nuclei was calculated.

### Cell growth experiments

Cell proliferation was assessed by seeding 2000 cells of ODin system cell lines or unmodified HCT116 cells 24h before bioPROTAC induction with 1 µg/mL doxycycline or treatment with 60 µM 10058-F4 MYC inhibitor ^48^ or DMSO (all reagents from Sigma Aldrich). Live cells were imaged every 2h for 72h on an Incucyte S3 system (Sartorius) using default settings. Cell confluence and average mCherry reporter fluorescence (used as a surrogate for bioPROTAC expression) were extracted as specified in the provided software (version 2019A). 72h end-point assays were performed using the CellTiter-Glo 2.0 cell viability assay (Promega) following manufacturer instructions, and measured in an EnVision plate reader. Statistical analysis was performed by an ordinary one-way ANOVA followed by Bonferroni’s multiple comparisons test (we assumed a normal distribution of the biological replicate values) using the GraphPad Prism 9 software. See analysis in Supplementary Table 6. Means were considered significantly different if the multiplicity-adjusted p-value < 0.05.

For EdU experiments, 50,000 A549 cells expressing bioPROTAC or controls were seeded in the presence or absence of 1 µg/mL doxycycline. Cells were incubated for 24 hours, and in the last 30 minutes of the incubation, in the presence of EdU. Cells were fixed and processed for IF using the Click-iT™ Plus EdU Proliferation kit (Alexafluor 647, Invitrogen). The rate of S-phase entry was measured by CellProfiler, and the average of 3 biological replicates was plotted and Two-way ANOVA statistical analysis was performed on GraphPad Prism 10.

For Crystal Violet experiments, 20,000 cells were seeded per well in clear 96-well plates in the presence or absence of doxycycline and incubated for 72 hours. Cells were then washed twice with PBS and fixed with 1% glutaraldehyde solution (Sigma) for 15 minutes followed by washes with PBS. Fixed cells were left to air dry overnight before staining with 0.1% Crystal Violet (Sigma) for 30 minutes. Wells were washed and left to air dry overnight before crystal violet extraction with 10% acetic acid for 30 minutes. Colourimetric measurements were taken at 590nm with a Clariostar plate reader. Each condition was performed in triplicate and normalised to the Crystal Violet intensity of the VH15^1KR^-L1-ICP0mt2 (with doxycycline); the average of three biological replicates was plotted and analysed by two-way ANOVA.

Spheroid formation assays were performed by seeding 5,000 cells per well in ultra-low attachment, round-bottom plates (Costar) followed by a brief centrifugation at 1,000rpm for 5 minutes. Cells were seeded in triplicate wells per day of CellTiter-Glo analysis. At days 1, 4, 6, and 8 from seeding, spheroids were transferred into white-walled 96-well plates (Thermoscientific) for incubation with Cell-Titer Glo reaction mixture. Luminescence data for each cell line was normalised to the day 8 readout for the -DOX DMSO condition.

### Immunoprecipitation

4×10^6^ HCT116 cells were transfected with 17 µg pFC14K encoding FLAG/Halo-tagged affinity proteins using FugeneHD as per vendor instructions. 2 days after transfection, cells were recovered and incubated 20 min in lysis buffer (1% NP40, 50 mM Tris HCl pH 7.5, 150 mM NaCl, 1 mM EGTA, 1 mM EDTA, 10 mM Glycerophosphate, 50 mM Sodium Fluoride, 0.27 M Sucrose, 5 mM Sodium Pyrophosphate, 1 mM Sodium orthovanadate) supplemented with cOmplete Mini EDTA-free protease inhibitor (Roche). Cell debris were removed by centrifugation and protein content quantified with the Pierce BCA Protein Assay Kit (Thermo Scientific) following vendor instructions. 12.5 µg cell lysate were diluted in reducing Laemmli sample buffer (Bio-Rad; reducing agent added as instructed) and incubated at 95°C for 5 min. 5 µL anti-FLAG M2 magnetic beads (Sigma) were pre-washed with lysis buffer and incubated overnight at 4°C with 250 µg cell lysate. Beads were subsequently washed 1x with lysis buffer and 2x with high salt wash buffer (1% NP40, 50 mM Tris HCl pH 7.5, 300 mM NaCl, 1 mM EGTA, 1 mM EDTA, 10 mM Glycerophosphate, 50 mM Sodium Fluoride, 0.27 M Sucrose, 5 mM Sodium Pyrophosphate, 1 mM Sodium orthovanadate) supplemented with protease inhibitors. Captured proteins were eluted in 1x reducing Laemmli sample buffer and analysed by immunoblotting.

### Immunoblotting

In mRNA transfection experiments, 3.2×10^5^ HCT116 cells were transfected with 1 µg mRNA using Lipofectamine RNAiMAX and harvested with 1x reducing Laemmli buffer after 17h. Where indicated, Lipofectamine RNAiMAX-mediated transfection of 20 nM Silencer Select CUL3 siRNAs (IDs s16048, s16049, s531239, s531241) or Silencer Select Negative Control No. 2 siRNA (all from Thermo Fisher) was performed 48h ahead of mRNA transfection. Constructs in ODin system cell lines were induced as previously specified. For all experiments, samples were prepared by lysing cells directly in 1x reducing Laemmli buffer, except in immunoprecipitation experiments (see above). SDS-PAGE and immunoblotting were performed using Criterion Cell and Criterion Blotter systems (Bio-Rad), as instructed by the manufacturer for Tris/Glycine buffers. PVDF membranes were incubated with blocking buffer (5% milk, 0.05% Tween in PBS) for 1h and overnight with one of the following antibodies: Anti-MYC Y69 (Abcam), anti-HA 16B12 (Enzo Life Sciences), anti-FLAG M2 (Sigma), anti-Vinculin E1E9V (Cell Signaling Technology, CST), anti-GAPDH 14C10 (Sigma), anti-MAX (Bethyl, cat. no. A302-866A-T) and anti-CUL3 (Bethyl, cat. no. A301-109A-T). Vinculin and GAPDH were used as loading controls (in the same blot as the remaining detected proteins). Washed membranes were incubated with the respective secondary antibodies conjugated to HRP (CST). Detection was performed using Immobilon Crescendo Western HRP substrate (Millipore) in a ChemiDoc imaging system (Bio-Rad) using the Image Lab 6.0.1 software (Bio-Rad). Where applicable, HRP was inactivated with 0.1% Sodium Azide (Sigma) and membranes re-incubated with a primary and secondary antibodies as instructed (usually in advance of anti-HA and anti-FLAG incubations). Band quantification by densitometry was performed using Image Lab, after which MYC integrated intensity values were normalized to those of vinculin and provided as a percentage relative to the mock control. Figure panels were constructed using Adobe Illustrator 2020 (version 24.2.1) and Adobe Photoshop CC 2019. Uncropped scans are shown in Supplementary Figure 9.

### Mass spectrometry and multiomics

Protein digests were prepared from cell pellets using S-Trap mini columns (ProtiFi). Cell pellets were resuspended in 1× S-Trap lysis buffer. Protein concentration was determined using a BCA assay kit (ThermoFisher) according to the user manual. After reduction and alkylation, equal amounts of protein were loaded onto S-Trap Mini columns and digested overnight. Equivalent amounts of resulting tryptic peptides were loaded onto Evotips and analysed on a Evosep One LC coupled to a timsTOF Pro2 or Ultra mass spectrometer (Bruker). The Spectra were searched using DIA-NN^49^ in a modified version of the quantms workflow^50^. An in-silico library was generated using human SwissProt sequences without isoforms from Uniprot release 2023_01 with carbamidomethylation as a fixed modification and N-term acetylation and methionine oxidation as variable modifications. Minimum peptide length was set at 7 and maximum 40 allowing up to 2 missed cleavages. The final outputs are filtered to 1% protein level and peptide level FDR, only considering uniquely mapping peptides. Differential expression analysis was performed using the MSstats R package^51^ and filtered to exclude proteins identified by a single peptide and features with less than three measurements. Data was median normalised at the peptide level and protein abundance was estimated via Tukey’s median polish from all peptides mapped to each target protein. Differential expression results were filtered to only include proteins with less than 1% adjusted p-value in the comparison for downstream analysis. For downstream analysis, changes of protein levels were deemed significant when Log_2_FC≤-0.5 or ≥0.5 and FDR-adjusted p-values (q-values)≤0.05. Pathway analysis was performed on Enrichr^52–54^ web interface using MSigDB Hallmark (2020).

### *In vitro* Rabbit Reticulocyte lysates

The *in vitro* translation assay was performed as previously described^27^ using the Flexi Rabbit reticulocyte lysate kit (Promega), using 1.5pmol of mRNA per reaction and reactions were run for 2 hours. Resulting proteins were processed for Western blot as above.

### RNA sequencing

A549 cells expressing VH15^1KR^-L1-ICP0^1KR^ or controls were seeded in 6-well plates (750,000 cells/well) and incubated for 24h in the presence or absence of doxycycline. Cells were washed in PBS and detached with Trypsin. Cells were resuspended in complete media and counted. 500,000 cells per condition were collected for RNA extraction for bulkRNA sequencing and processed using Monarch Total RNA Miniprep kit (NEB). Library preparation, sequencing and bioinformatic analysis was performed by Genewiz by Azenta. Illumina Sequencing was performed on a NovaSeq 2×150 and sequence reads were aligned to the GRCh38 (GRCh38.p7, NCBI:GCA_000001405.22). For downstream analysis, gene expression changes were deemed significant when Log_2_FC≤-0.5 or ≥0.5 and FDR-adjusted p-values (q-values)≤0.05. Pathway analysis was performed on Enrichr^52–54^ web interface using MSigDB Hallmark (2020).

### Ethical approval and Animal studies

Animal experiments received ethical approval and were conducted in accordance with The Home Office UK guidelines, under project licence PP9816261 (Walid Khaled), which was evaluated and approved by the Animal Welfare and Ethical Review Body at the University of Cambridge. *NU(NCr)-Foxn1nu* mice were obtained from Charles River. All animals were maintained under SPF conditions, fed a regular diet in a pathogen-free facility, and housed on a 12-hour light/dark cycle with continuous access to food and water. Prior to injection, asynchronous growing cells were confirmed to be Mycolasma-free, harvested and mixed 1:1 with Cultrex RGF BME (Biotechne). Subcutaneous injections were used to generate xenografts in the flanks of mice, with 500,000 cells injected per flank. Schedule 1 method of euthanasia was used for all mice. Animals were killed and subcutaneous tumours, lung liver and spleen were collected and fixed in 10% neutral-buffered formalin overnight at room temperature, before moving tissues to 70% ethanol and processed into paraffin wax.

### Histology

Tissues were sectioned at 4 µm and mounted on slides (FisherScientific). For each stain, sections were rehydrated, blocked with 2.5% horse serum, 1% BSA in PBS (blocking buffer) and stained with primary antibodies in blocking buffer overnight at 4°C. The following day, tissue sections were washed in PBS, stained with secondary antibodies for 1 hour at room temperature and with Hoechst 33258 (Sigma-Aldrich) diluted 1:5000 in PBS for 5 minutes. Coverslips were mounted using ProLong Gold mounting media (ThermoFisher). Each slide was then imaged with a Leica DM6 B using LASX software. Primary antibodies: Goat anti-GFP polyclonal antibody (ab5450, abcam), Rabbit anti-RFP polyclonal (600-401-379, Rockland), Rabbit anti-HA-Tag monoclonal antibody (Cell Signalling Technologies, 3724S), Rabbit anti-Cleaved Caspase 3 monoclonal antibody (Cell Signalling Technologies, 9664S). Secondary antibodies: Donkey anti-Goat IgG (H+L) Cross-Adsorbed Secondary Antibody, Alexa Fluor 488 (A11055), Donkey anti-Rabbit IgG (H+L) Cross-Adsorbed Secondary Antibody, Alexa Fluor 555 (A31572). Wheat germ agglutinin (WGA647, Invitrogen) staining was performed concomitantly to secondary antibody incubation. We measured signal areas in an automatic and unbiased manner using a custom script for Fiji^55^. The script segments tissue area using WGA labelling, and detects GFP-positive regions within the tissue. Errors due to variability in intensity distribution between samples were reduced by using a minimum coefficient of variation of 0.3 to identify images containing a signal as well as background distribution. These selected images were thresholded independently using Otsu’s method^56^ to map signal areas. For quantification of CC3 expression, quantification of GFP- and CC3-positive areas was performed on 4 random tiles per tumour, with selected images thresholded independently using Otsu’s method based on areas of positive signal.

