## Supplemental data for "Combinatorial screening of biological degrader modules identifies heterobifunctional molecules that efficiently destroy MYC, leading to rapid tumour cell death *in vitro* and *in vivo*"

**Supplemental Figures 1 to 10**

**Supplemental Tables 1 to 9 (additional files)**

**Supplementary Figures:**

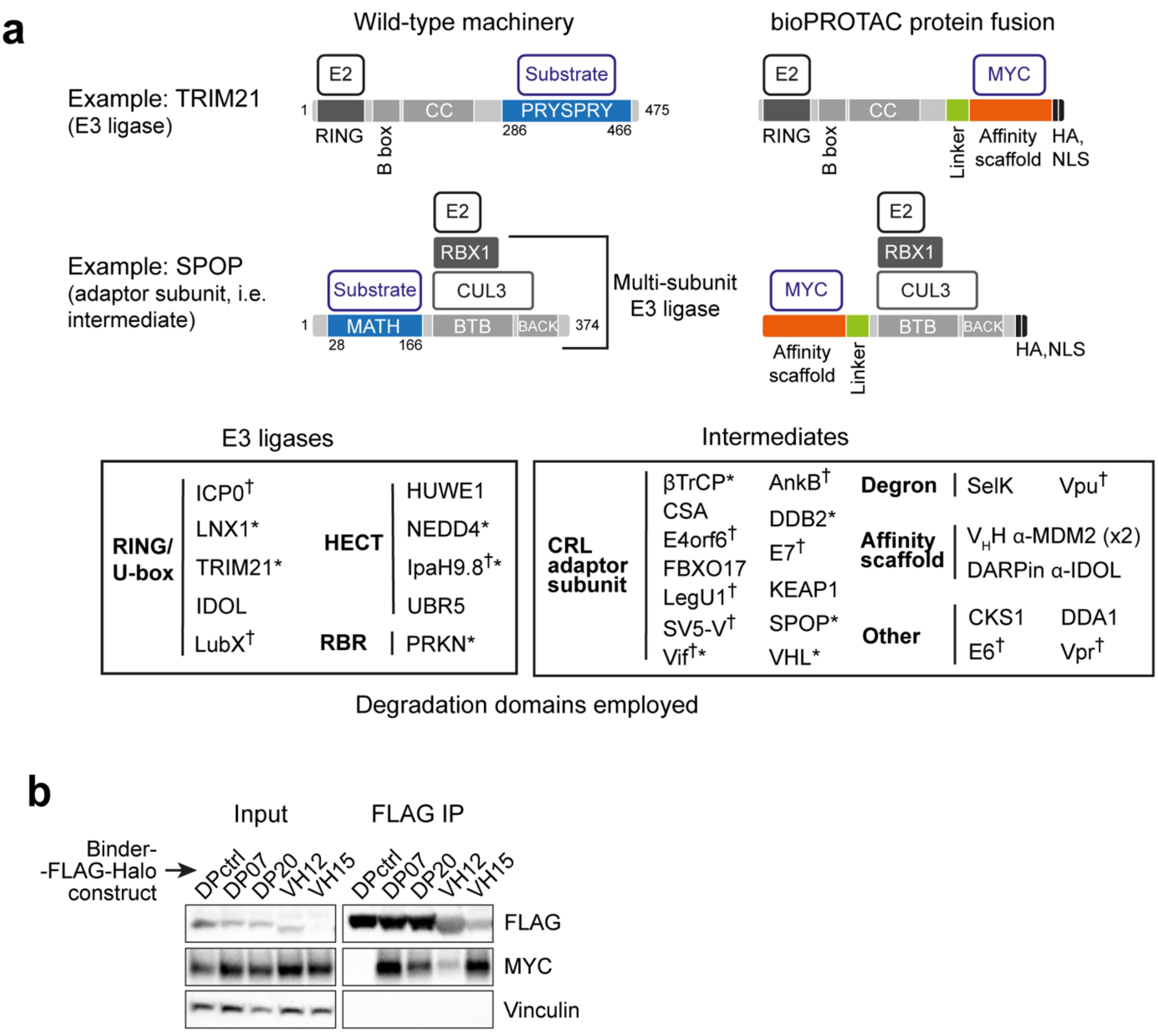


Supplementary Figure 1

**Supplementary Figure 1: Component selection and design of MYC-targeting bioPROTACs.** (**a**) Above: bioPROTACs are constructed by extracting functional domains from E3 ligases or E3 ligase-recruiting proteins, which are fused to a MYC-recruiting scaffold. The E3 remnant (“degradation domain”, DD) must retain the capacity to recruit loaded E2 ubiquitin-conjugating enzymes (E2~Ub) to facilitate the transfer of ubiquitin to the target. Two specific engineering examples are given for TRIM21[11] and SPOP[6]-based bioPROTACs (an E3 ligase and intermediate, respectively). Below: Unique proteins employed in this study as DDs, categorized by E3 ligase family or intermediate type. Note that the DD toolbox may include different truncations or orientations of the indicated proteins. Proteins marked as “other” don’t usually recruit a substrate on their own and are either CRL interactors which do not act independently of substrate adaptors (DDA1, CKS1) or pathogenic proteins which modulate an E3 ligases’ specificity (Vpr, E6). Each bioPROTAC contains an HA tag for detection and an NLS for co-localization with MYC. (**b**) FLAG immunoprecipitation (IP) experiments following transient transfection of plasmids encoding for the indicated FLAG-Halo-tagged MYC binders or nonbinding DARPin (DPctrl). Input is 5% of cell lysate. Vinculin is used as a control for loading and purity. Abbreviations: CRL, Cullin-

RING Ligase; HECT, Homologous to E6AP C-terminus; RBR, RING-between-RING; RING, Really Interesting New Gene; Ub, Ubiquitin. *Used in the literature in similar approach (two DD designs exist for βTrCP). †domain is of pathogenic origin.

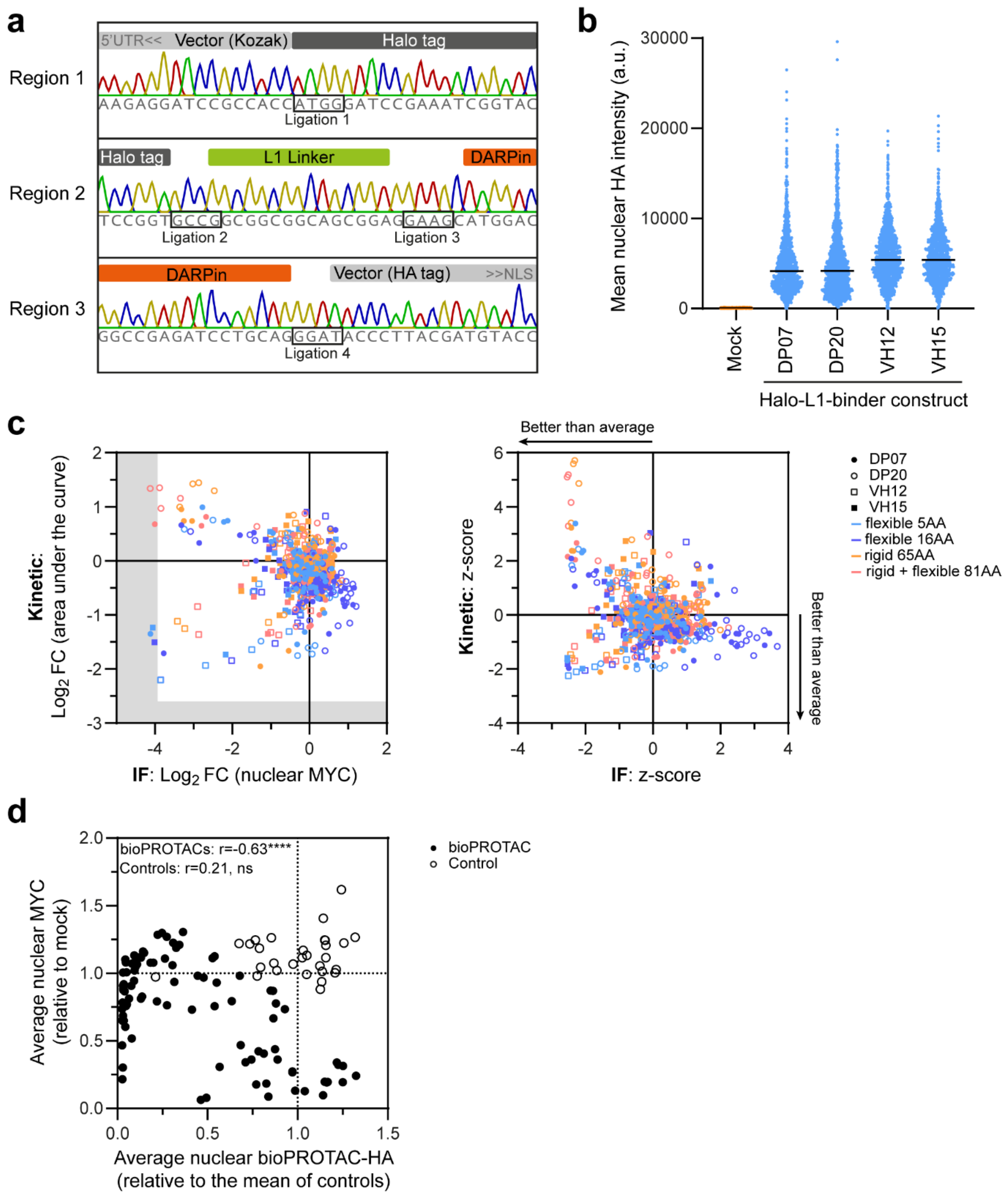


Supplementary Figure 2

**Supplementary Figure 2: Method establishment and usage for screening of DP07, DP20, VH12 and VH15-containing anti-MYC bioPROTACs.** Related to Fig. 1c. (**a**) Sanger sequencing chromatogram snapshots covering the four joining regions of a Halo-L1-DARPin control construct assembled with the automated workflow (example of N=2). (**b**) Measured bioPROTAC-HA expression in single cells by IF for Halo-L1 controls in the MYC bioPROTAC screening (example of N=3). The entire cell population is included, and black lines indicate the median abundance. a.u., arbitrary units. (**c**) Nuclear MYC abundance (IF) and area under the curve (kinetic assay) fold changes (FC) in $\log_2$ scale (left) or as z-scores (right) in the primary screening (N=1) for (DP/VH discovery). bioPROTACs were delivered as mRNA in HCT116 cells, and FCs are relative to controls where the degradation domain is replaced by a Halo tag. Symbols designate the MYC binder: DP07 (full circle), DP20 (empty circle), VH12 (full square) and VH15 (empty square), whereas colours identify the linker: flexible 5AA (light blue), flexible 16AA (royal blue), rigid 65AA (orange), rigid+flexible 81AA (coral). Shaded areas depict the detection limit (IF) or most extensive realistic MYC downregulation (kinetic assay), determined by 100 μg/mL cycloheximide treatment. (**d**) MYC abundance effect dependence on construct expression (normalized to mock and the average of controls, respectively), measured by IF in the confirmatory screening (average of N=3) when HA-tagged bioPROTACs (full circle) or controls (empty circle) are expressed. Pearson correlation tests were performed and the resulting correlation coefficient (r) are indicated. p-value: ****<0.0001, ns=non-significant (>0.05).

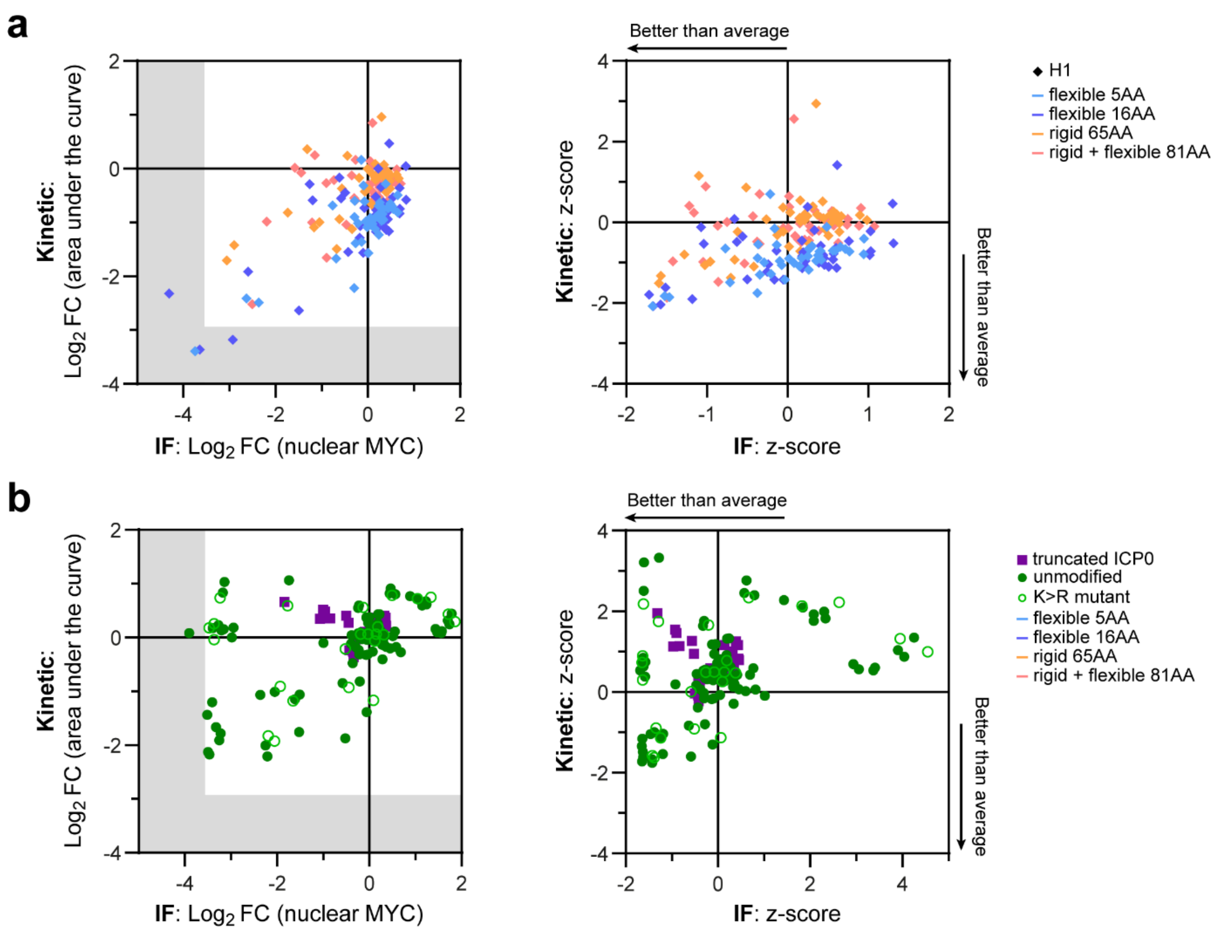


Supplementary Figure 3

**Supplementary Figure 3: H1 discovery and hit validation & optimization primary screening results.** Related to Fig. 1d. (**a**, **b**) Fold change (FC) in Log$_2$ scale (left) or z-score (right) correlation between MYC abundance assay outputs for the follow-up primary MYC bioPROTAC screening (N=1) in HCT116 cells transfected with mRNA. (**a**) H1 discovery primary screening data, whereby FCs are relative to controls lacking a functional degradation domain. Datapoints refer to expression of H1-containing bioPROTACs with flexible 5AA (light blue), flexible 16AA (royal blue), rigid 65AA (orange), rigid+flexible 81AA (coral) linkers. (**b**) Results from the optimization & validation screening subset (primary screening), whereby FCs are calculated vs controls unable to recruit MYC. Unmodified (green full circle), K>R mutant (green empty circle) or bioPROTACs containing further ICP0 truncations (purple square) were expressed. Shaded areas depict MYC downregulation by cycloheximide, taken as the background signal.

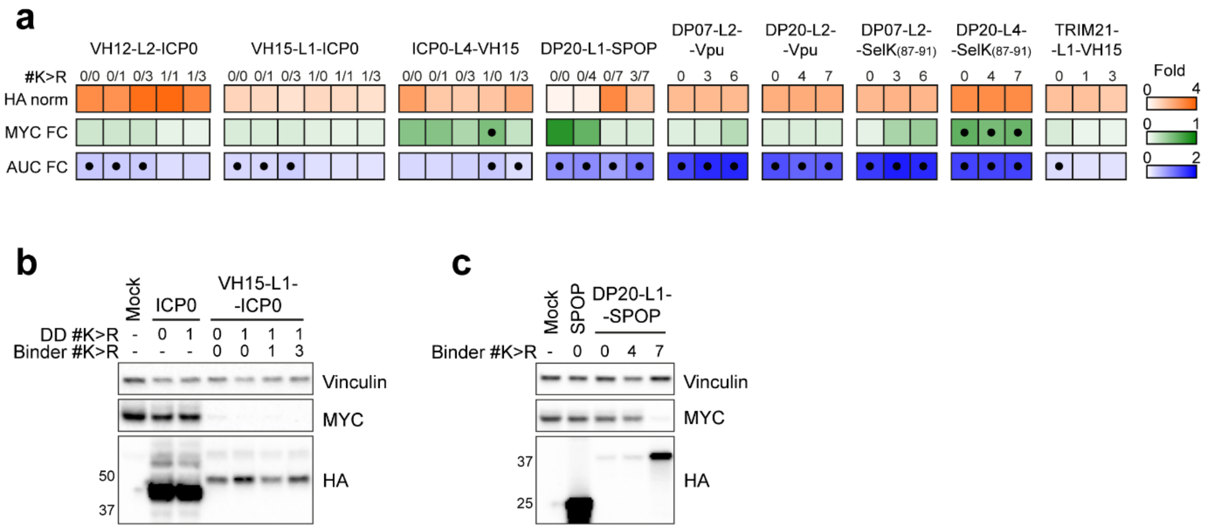


Supplementary Figure 4

**Supplementary Figure 4: KR substitutions in bioPROTAC abundance and effect.** (**a**) Heatmap of validation & optimization confirmatory screening constructs (values are the average of N=3), depicting nuclear bioPROTAC-HA intensity normalized to mean of controls (“HA”), nuclear MYC FC in IF (“MYC FC”) and the area under the curve FC from the kinetic assay (“AUC FC”). Black dots highlight nonsignificant values (q-value<0.05 in FDR-corrected t-tests). #K>R: number of Lys to Arg substitutions. Numbers in x/y format represent KR substitutions in the degradation domain and in the target-binding domain, respectively, and a single number means that all mutations are in the target-binding domain. (**b**, **c**) Immunoblotting following HCT116 mRNA-based expression of bioPROTACs bearing the indicated number of K>R substitutions in the ICP0 degradation domain (DD) or binders (VH15 or DP20). Note that for immunoblotting experiments, screening workflow simplifications were not employed, and the same mRNA amount was transfected in all samples. A representative example of three independent experiments is shown. Numbers left of HA blots indicate the molecular weight of protein markers. Vinculin is used as loading control and anti-HA detects bioPROTACs.

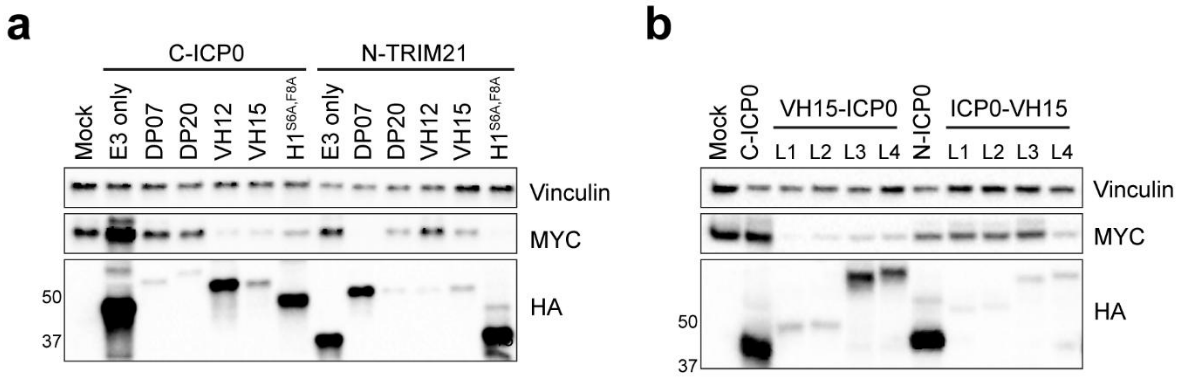


Supplementary Figure 5

**Supplementary Figure 5: Composition preferences for selected MYC-targeting bioPROTACs.** Immunoblotting following HCT116 mRNA-based expression of bioPROTACs (HA-tagged) with varying compositions to investigate (**a**) DD-binder relationships (constructs have the flexible 16AA linker) and (**b**) ICP0 positioning and the contribution of the different linkers for MYC downregulation. For these experiments, mRNA was re-synthesized without the screening workflow simplifications and the same amount transfected in all samples. A representative example of three independent experiments is shown. Linkers: L1, flexible 5AA; L2, flexible 16AA; L3, rigid 65AA and L4, rigid+flexible 81AA. Numbers left of HA blots indicate the molecular weight of protein markers (note that the H1 peptide is not detected due to its small size). Vinculin is used as loading control and anti-HA detects bioPROTACs.

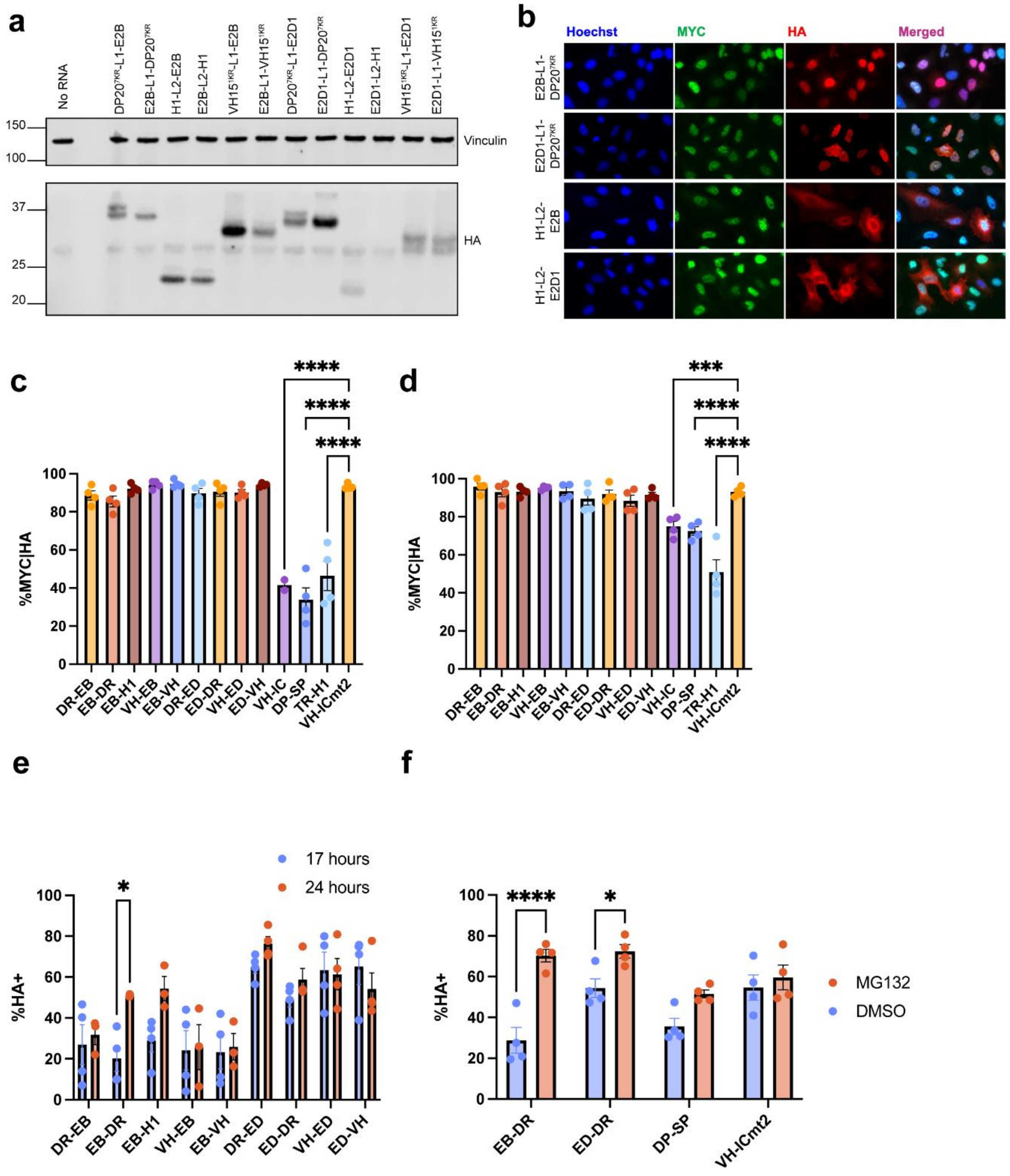


Supplementary Figure 6

**Supplementary Figure 6: Screen of E2-based bioPROTACs against MYC.** (**a**) Representative of n=3 Western blot of protein resulting from the *in vitro* translation assay rabbit reticulocyte lysate supplemented with No RNA or mRNA encoding E2-based bioPROTACs (HA). Vinculin was used as a loading control (**b**) Representative images of E2-based bioPROTACs (HA) showing correct nuclear localisation (E2B-L1-DP20$^{7KR}$ and E2D1-L1-DP20$^{7KR}$) and displaying incorrect, cytoplasmic localisation (H1-L2-E2B, H1-L2-E2D1). (**c, d**) Quantification of the immunofluorescence screen in A549 for E2-based constructs, E3-based bioPROTACs, and non-degrader control. Data shows the percentage of MYC-positive HA-positive nuclei of n=4 biological replicates ± SEM, analysed by one-way ANOVA at 17 hours (c) and 24 hours (d) post-transfection. (**e**) Quantification of HA-positive nuclei at 17 and 24 hours post-transfection for A549 cells expressing E2-based bioPROTACs. (**f**) Quantification of HA-positive nuclei of A549 cells transfected with E2B-L1-DP20$^{7KR}$, E2D1-L1-DP20$^{7KR}$, DP20$^{7KR}$ -L1-SPOP or non-degrader control VH15$^{1KR}$-L1-ICP0mt2 for 24 hour and treated with DMSO or proteasome inhibitor MG132 (10 μM) in the last hour prior to fixation. p-values: *<0.05, ***<0.001, ****<0.0001.

Sample Key: DR-EB= DP20$^{7KR}$-L1-E2B; EB-DR=E2B-L1-DP20$^{7KR}$; E2B-H1=EB-L2-H1; VH-EB= VH15$^{1KR}$-L1-E2B; EB-VH=E2B-L1-VH15$^{1KR}$; DR-ED=DP20$^{7KR}$-L1-E2D1; ED-DR=E2D1-L1-DP20$^{7KR}$; VH-ED=VH15$^{1KR}$-L1-E2D1; ED-VH=E2D1-L1-VH15$^{1KR}$; VH-IC=VH15$^{1KR}$-L1-ICP0$^{1KR}$; DP-SP=DP20$^{7KR}$-L1-SPOP; TR-H1=TRIM21-L2-H1; VH-ICmt2=VH15$^{1KR}$-L1-ICP0mt2

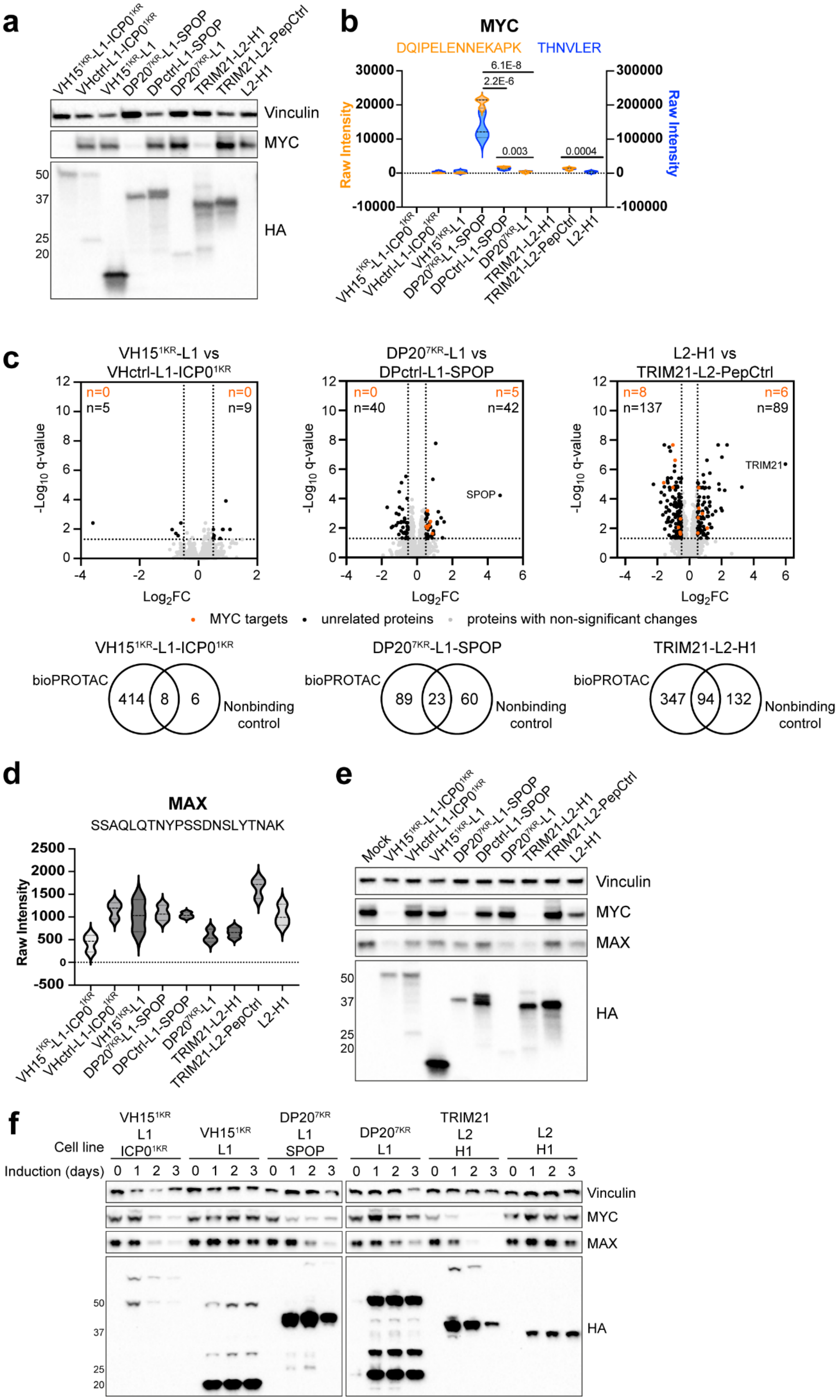


Supplementary Figure 7

**Supplementary Figure 7: Analysis of proteome changes by mass spectrometry discriminates on-/off-target effects of anti-MYC bioPROTACs.** Related to Figure 2c. (**a**) Validation of mRNA-based bioPROTAC-HA expression and MYC downregulation in the HCT116 cell lysates analysed by mass spectrometry (one example of n=3). Controls include VHctrl, DPctrl and PepCtrl as non-MYC binding moieties or lack a degradation domain. Numbers left of the HA blot indicate the molecular weight of protein markers. Vinculin is used as loading control, and anti-HA detects bioPROTACs. (**b**) Raw intensities of MYC peptides (DQIPELENNEKAPK and THNVLER, orange and blue, respectively) and adjusted p-values shown as analysed by mass spectrometry. (**c**) Volcano plots depicting proteome changes upon expression of the indicated constructs (data is from 3 independent experiments). Significant changes of protein levels ($Log_2FC \leq -0.5$ or $\geq 0.5$ and FDR-adjusted p-values (q-values)$\leq 0.001$) are depicted for MYC transcriptional targets regulated by the binders (orange), as binders may themselves inhibit MYC, or for unrelated proteins (black); (grey) represents proteins with non-significant changes in levels. Venn diagrams portraying the number of downregulated proteins found upon expression of the indicated bioPROTACs (from Fig. 2c) and in the nonbinding controls (both compared to unfunctionalized binder control) ($Log_2FC \leq -0.5$ or $\geq 0.5$ and FDR-adjusted p-values (q-values)$\leq 0.05$). Elements common to both datasets are likely off-target proteins degraded by the bioPROTAC's degradation domain. (**d**) Raw intensities of MAX peptides (SSAQLQTNYPSSDNSLYTNAK) analysed by mass spectrometry. (**e**) Immunoblotting after expression of the indicated constructs by mRNA transfection for 17h. (**f**) Similar analysis following induction of bioPROTAC-T2A-mCherry constructs with 1 µg/mL doxycycline in stable cell lines for the indicated time-points. Representative data of 3 independent experiments are shown. Numbers left of HA blots indicate the molecular weight of protein markers. Ectopically-expressed constructs were detected with anti-HA tag antibody and vinculin was used as loading control.

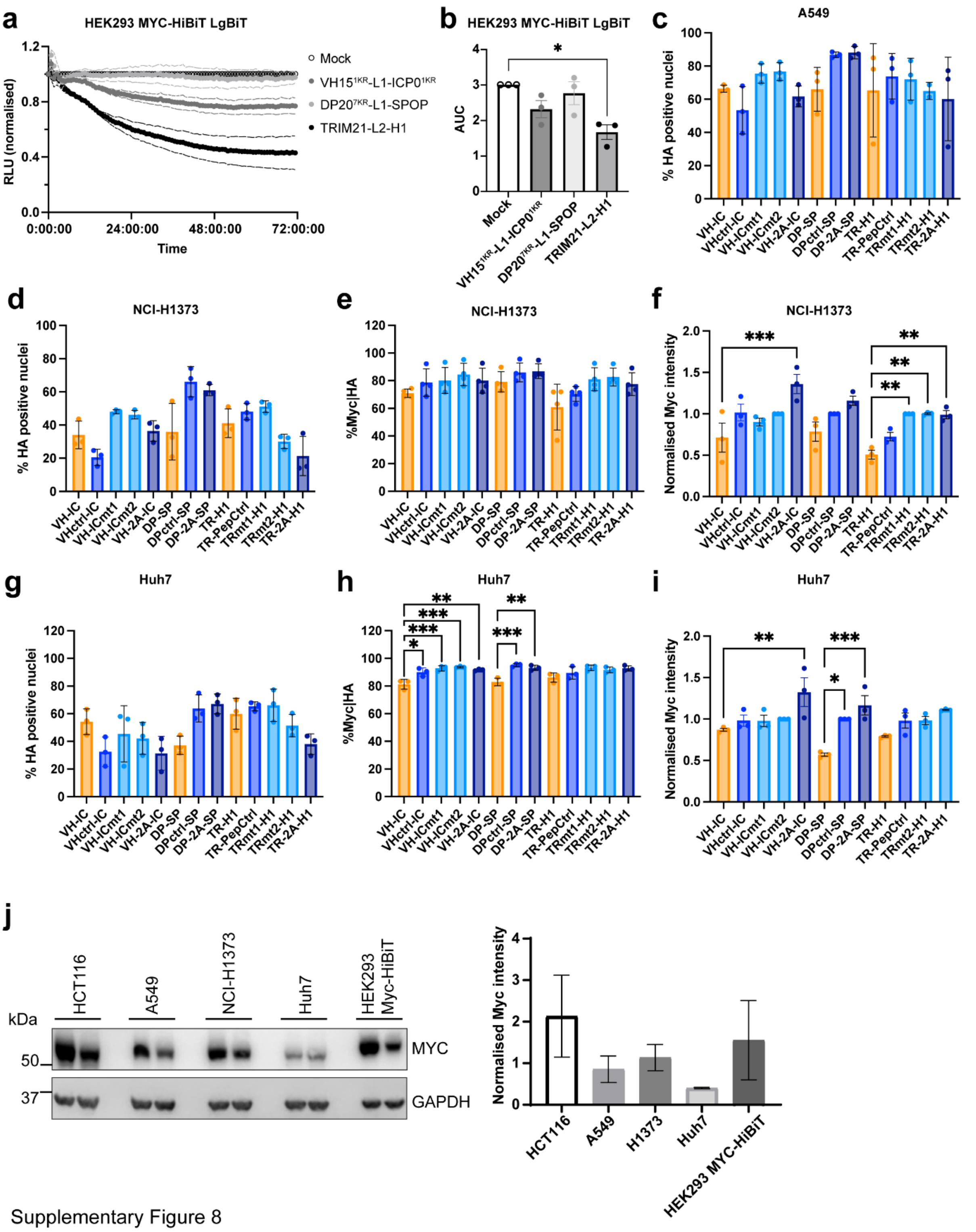


Supplementary Figure 8

**Supplementary Figure 8: Lead bioPROTACs degrade MYC in multiple cell lines** (**a**) Time-course data depicting normalized relative light units (RLU) in HEK293 MYC-HiBiT, LgBiT cells transfected with bioPROTACs or mock-transfected. Data is the mean (solid lines) ±standard error of the mean (SEM, dotted lines) of n=3 biological replicates. (**b**) Area under the curve (AUC) analysis of n=3 biological replicates of time-course HEK293 MYC-HiBiT, LgBiT cells as in (a). Kruskal-Wallis test with multiple comparisons to mock was performed. Transfection efficiency of A549 (**c**, related to Figure 2), NCI-H1373 (**d**), and Huh-7 (**g**) cells transfected with bioPROTACs or controls (HA-tagged) was scored as a percentage of HA-positive nuclei following immunofluorescence staining 17 hours post-transfection.

MYC degradation was assessed by immunofluorescence by scoring the percentage of transfected (HA positive) nuclei positive for MYC (**e**, **h**) or by measuring the per-transfected nuclei MYC intensity, shown normalised to a non-functional control, per construct (**f**, **i**). Data is the percentage of MYC-positive HA-positive nuclei of at least n=3 biological replicates ± SEM, analysed by One-Way ANOVA. (**j)** Endogenous MYC levels in all cell lines used were compared by Western blot. Data shown as mean ± SD of n=2 biological replicates as MYC intensity normalised to relative GAPDH intensity. *$p<0.05$, ** $p<0.005$, *** $0.0001<p<0.0005$, **** $p<0.0001$. Sample key: VH-IC=VH15$^{1KR}$-L1-ICP0$^{1KR}$; VHctrl-IC= VHctrl-L1-ICP0$^{1KR}$; VH-ICmt1=VH15$^{1KR}$-L1-ICP0mt1; VH-ICmt2= VH15$^{1KR}$-L1-ICP0mt2; VH-2A-IC= VH15$^{1KR}$-T2A-ICP0$^{1KR}$; DP-SP=DP20$^{7KR}$-L1-SPOP; DPctrl-SP=DPCtrl-L1-SPOP; DP-2A-SP= DP20$^{7KR}$-T2A-SPOP; TR-H1=TRIM21-L2-H1; TR-PepCtrl=TRIM21-L2-PeptideCtrl; TRmt1-H1=TRIM21mt1-L2-H1; TRmt2-H1=TRIM21mt2-L2-H1; TR-2A-H1= TRIM21-T2A-H1

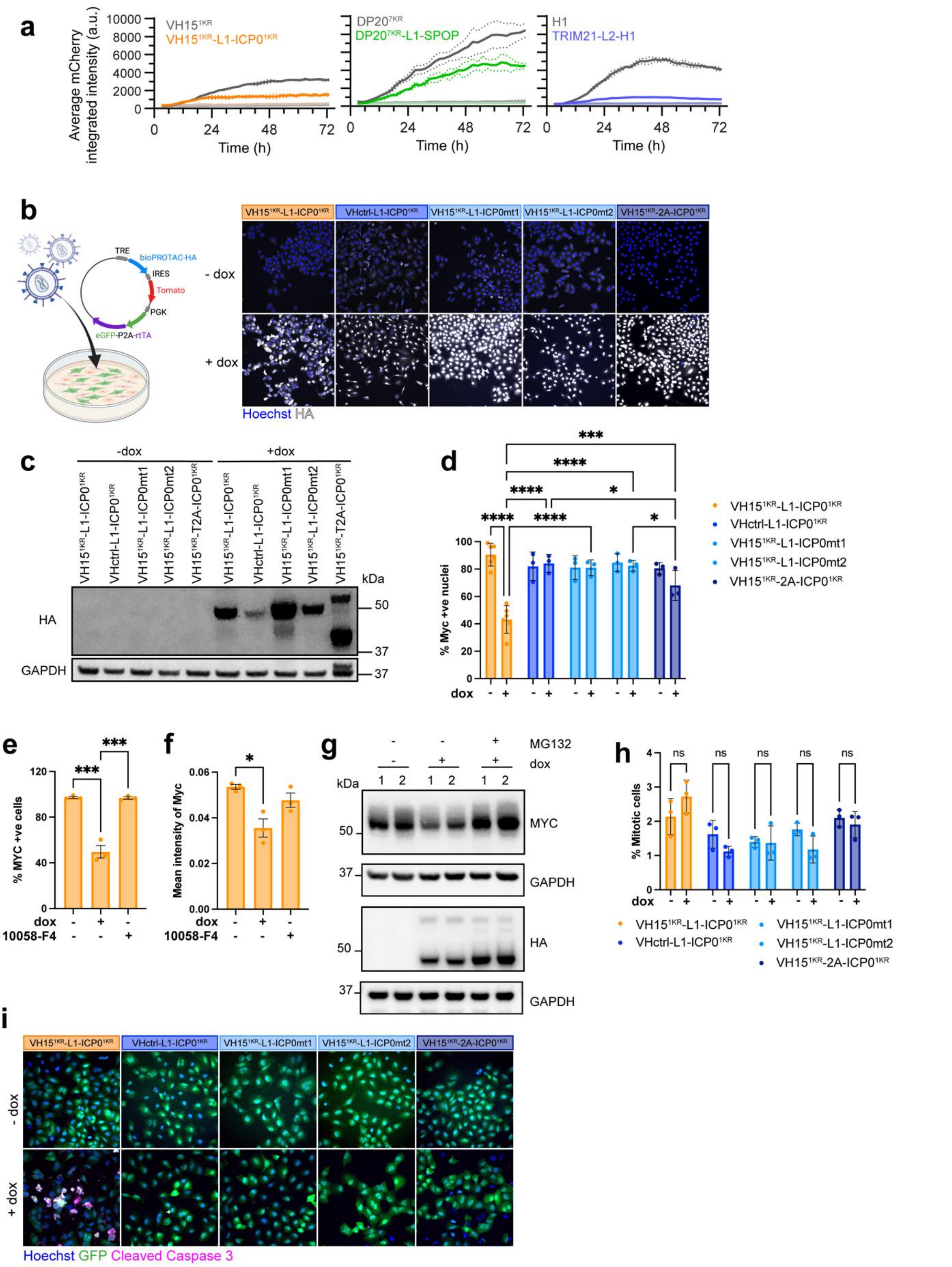

Supplementary Figure 9

**Supplementary Figure 9: Validation of stable cell lines expressing bioPROTAC or controls.** Related to Figures 3 and 4a (**a**) mCherry reporter expression (average integrated intensity per cell) of the indicated HCT116 doxycycline-inducible cell lines expressing construct-T2A-mCherry fusions. Strong colours designate induction of the indicated construct, as opposed to non-induced conditions represented by fainter colours. Continuous lines are the mean of 3 technical replicates for a representative biological replicate (of n=3). Dotted lines depict the error (SD). (**b**) Graphical representation of lentivirally transduced A549 expressing constitutively GFP-P2A-rtTA and conditional expression of bioPROTAC-HA (or control) and Tomato under the control of the tetracycline response element. Representative images of single-cell clones expressing VH15$^{1KR}$-L1-ICP0$^{1KR}$ bioPROTAC or controls in the presence of doxycycline (24 hours). (**c**) Western blot of A549 stable cell lines (single-cell clones) expressing VH15$^{1KR}$-L1-ICP0$^{1KR}$ bioPROTAC or controls following incubation with doxycycline for 24 hours. Validation of MYC degradation in A549 stable cell lines following 24-hour induction with doxycycline (**d**) or 10058-F4 treatment (50 μM) (**e**). Data shows the percentage of MYC-positive nuclei of at least n=3 biological replicates ± SEM, analysed by two-way ANOVA (d) or ordinary one-way ANOVA (e) with multiple comparisons. (**f**) Quantification of mean nuclear MYC intensity in VH15$^{1KR}$-L1-ICP0$^{1KR}$-encoding cells, as assessed by immunofluorescence, following 24-hour incubation with doxycycline or 10058-F4. Data is shown as mean ± SEM (of n=3) and analysed by ordinary one-way ANOVA with multiple comparisons. (**g**) Validation of proteasomal degradation of MYC in A549 stable cell line expressing VH15$^{1KR}$-L1-ICP0$^{1KR}$ treated with or without doxycycline for 24 hours or with the addition of proteasomal inhibitor MG132 (10 μM) for 2 hours. (**h**) Representative images of cleaved caspase 3 (CC3) immunofluorescence staining of cells expressing VH15$^{1KR}$-L1-ICP0$^{1KR}$ bioPROTAC or controls in the presence of doxycycline (24 hours). p-values: *<0.05, ***<0.001, ****<0.0001.

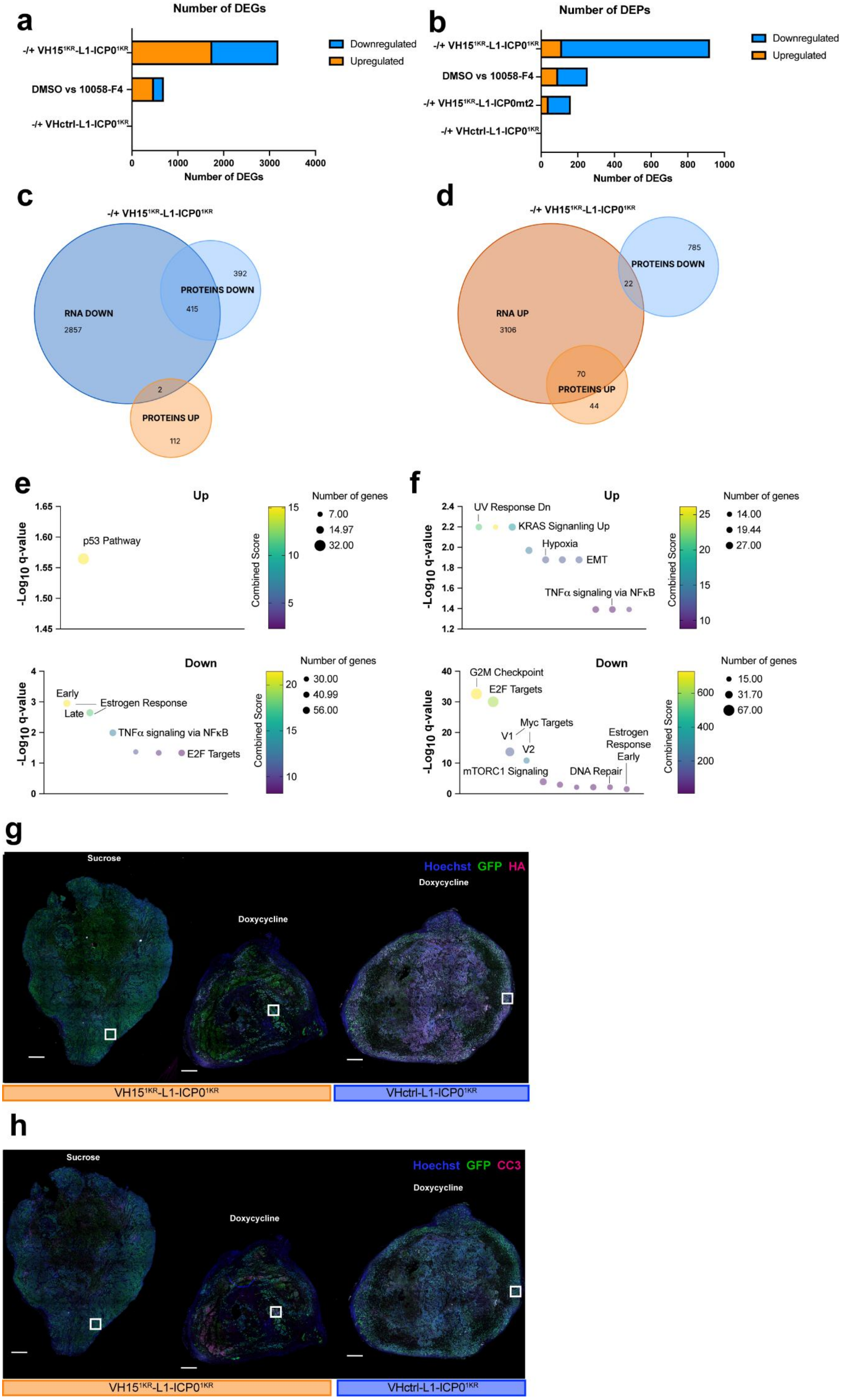
a
Number of DEGs
-/+ VH15$^{1KR}$-L1-ICP0$^{1KR}$
DMSO vs 10058-F4
-/+ VHctrl-L1-ICP0$^{1KR}$
0
1000
2000
3000
4000
Number of DEGs
Downregulated
Upregulated
b
Number of DEPs
-/+ VH15$^{1KR}$-L1-ICP0$^{1KR}$
DMSO vs 10058-F4
-/+ VH15$^{1KR}$-L1-ICP0mt2
-/+ VHctrl-L1-ICP0$^{1KR}$
0
200
400
600
800
1000
Number of DEGs
Downregulated
Upregulated
c
-/+ VH15$^{1KR}$-L1-ICP0$^{1KR}$
RNA DOWN
2857
PROTEINS DOWN
392
415
2
PROTEINS UP
112
d
-/+ VH15$^{1KR}$-L1-ICP0$^{1KR}$
785
PROTEINS DOWN
22
RNA UP
3106
70
PROTEINS UP
44
e
Up
-Log10 q-value
1.65
1.60
1.55
1.50
1.45
p53 Pathway
Combined Score
15
10
5
Number of genes
7.00
14.97
32.00
Down
-Log10 q-value
4
3
2
1
0
Early
Late
Estrogen Response
TNFα signaling via NFκB
E2F Targets
Combined Score
20
15
10
Number of genes
30.00
40.99
56.00
f
Up
-Log10 q-value
2.4
2.2
2.0
1.8
1.6
1.4
1.2
UV Response Dn
KRAS Signanling Up
Hypoxia
EMT
TNFα signaling via NFκB
Combined Score
25
20
15
10
Number of genes
14.00
19.44
27.00
Down
-Log10 q-value
40
30
20
10
0
G2M Checkpoint
E2F Targets
Myc Targets
V1
V2
mTORC1 Signaling
Estrogen Response Early
DNA Repair
Combined Score
600
400
200
Number of genes
15.00
31.70
67.00
g
Sucrose
Doxycycline
Doxycycline
Hoechst GFP HA
VH15$^{1KR}$-L1-ICP0$^{1KR}$
VHctrl-L1-ICP0$^{1KR}$
h
Sucrose
Doxycycline
Doxycycline
Hoechst GFP CC3
VH15$^{1KR}$-L1-ICP0$^{1KR}$
VHctrl-L1-ICP0$^{1KR}$

**Supplementary Figure 10: BulkRNA sequencing of A549 cells expressing bioPROTACs or controls and histological analysis of xenograft tumours.** Related to Figure 4. (**a**) Total number of Differentially expressed genes (DEGs) per analysed comparison. (**b**) Total number of Differentially expressed proteins (DEPs) per analysed comparison. (**c-d**) Overlap of DEGs and DEPs recorded from VH15$^{1KR}$-L1-ICP0$^{1K}$. (**e-f**) Pathway Analysis of MSigDB Hallmark genes of significant DEGs for: VH15$^{1KR}$-L1-ICP0$^{1KR}$ with and without doxycycline (**e**); or MYC inhibition with 10058-F4 vs DMSO (**f**). Significant changes of expression levels ($Log_2FC \leq -0.5$ or $\geq 0.5$ and FDR-adjusted p-values (q-values)$\leq 0.001$) from 3 independent experiments. (**g-h**) Histological scans of A549 xenograft tumours from Athymic nude mice of VH15$^{1KR}$-L1-ICP0$^{1KR}$, +/- doxycycline or non-binder control VH1ctrl-L1-ICP0$^{1KR}$. Immunofluorescence for HA (**g**) or Cleaved Caspase 3 (**h**). White boxed areas shown in Figure 4 (0.5 mm scale bar).
